# A ubiquitin-proteasome pathway degrades the inner nuclear membrane protein Bqt4 to maintain nuclear membrane homeostasis

**DOI:** 10.1101/2022.12.29.522265

**Authors:** Toan Khanh Le, Yasuhiro Hirano, Haruhiko Asakawa, Koji Okamoto, Tatsuo Fukagawa, Tokuko Haraguchi, Yasushi Hiraoka

**Affiliations:** Nuclear Dynamics Group, Graduate School of Frontier Biosciences, Osaka University, Suita 565-0871, Japan; Laboratory of Mitochondrial Dynamics, Graduate School of Frontier Biosciences, Osaka University, Suita 565-0871, Japan; Laboratory of Chromosome Biology, Graduate School of Frontier Biosciences, Osaka University, Suita 565-0871, Japan

**Keywords:** Inner nuclear membrane, Protein degradation, Fission Yeast, Bqt3, Bqt4

## Abstract

Aberrant accumulation of inner nuclear membrane (INM) proteins is associated with deformed nuclear morphology and mammalian diseases. However, the mechanisms underlying the maintenance of INM homeostasis remain poorly understood. In this study, we explored the degradation mechanisms of the INM protein, Bqt4, in the fission yeast *Schizosaccharomyces pombe*. We have previously shown that Bqt4 interacts with the transmembrane protein Bqt3 at the INM and is degraded in the absence of Bqt3. Here, we revealed that excess Bqt4, unassociated with Bqt3, was targeted for degradation by the ubiquitin-proteasome system localized in the nucleus and Bqt3 antagonized this process. The degradation process involved the Doa10 E3 ligase complex at the INM. Bqt4 is a tail-anchored protein and extraction from the membrane by the Cdc48 complex is required for its degradation. The C-terminal transmembrane domain of Bqt4 is necessary and sufficient for proteasome-dependent protein degradation. Accumulation of Bqt4 at the INM impaired cell viability with nuclear envelope deformation, suggesting that quantity control of Bqt4 plays an important role in nuclear membrane homeostasis.

Summary statement

Appropriate levels of the inner nuclear membrane protein Bqt4 are maintained by ubiquitin-proteasome-mediated degradation. Aberrant accumulation of Bqt4 disturbs nuclear membrane homeostasis.

## Introduction

The eukaryotic nucleus is enclosed by the nuclear envelope (NE), which separates the nucleoplasm from the cytoplasm. The NE comprises inner and outer nuclear membranes (INM and ONM, respectively). The ONM is continuous with the endoplasmic reticulum (ER). The INM is involved in the organization and regulation of chromosomes by modulating chromatin potential for gene expression, DNA replication, DNA damage repair, and genome stability (Mekhail and Moazed, 2010; Hirano et al., 2020a; Pawar and Kutay, 2021). Excess amounts of INM proteins result in their aberrant accumulation at the NE, often causing nuclear deformation and malfunctioning (Gonzalez et al., 2012). The aberrant accumulation of INM proteins has been linked to certain mammalian diseases. For example, accumulation of the INM protein SUN1 was found to be pathogenic in human laminopathies such as Emery-Dreifuss muscular dystrophy and Hutchinson-Gilford progeria syndrome (Chen et al., 2012). Therefore, the amount of INM proteins must be regulated. However, the mechanisms that regulate levels of resident INM proteins to maintain INM homeostasis remain poorly understood.

Mechanisms for ER membrane homeostasis have been well studied: ER membrane proteins are regulated through the ER-associated degradation (ERAD) system, which is involved in degrading misfolded or mislocalized ER proteins as a means of protein quality control (Vembar and Brodsky, 2008; Ruggiano et al., 2014; Christianson and Carvalho, 2022). ERAD involves the ubiquitin-proteasome system (UPS), in which polyubiquitin chains are covalently attached to a substrate protein, leading to its recognition and degradation by the 26S proteasome complex. Ubiquitination reactions involve three different classes of enzymes: E1 ubiquitin-activating enzymes, E2 ubiquitin-conjugating enzymes, and E3 ubiquitin ligases, such as Hrd1 and Doa10 (human MARCHF6/TEB4). INM-associated degradation (INMAD) has been proposed as an extension of the ERAD to maintain INM homeostasis, which also depends on the UPS (Foresti et al., 2014; Khmelinskii et al., 2014; Koch and Yu, 2019; Smoyer and Jaspersen, 2019; Mannino and Lusk, 2022). It has been shown that in the budding yeast *Saccharomyces cerevisiae*, ERAD and INMAD require additional machinery for the degradation of integral membrane proteins, involving Cdc48/p97, an ATPase associated with diverse cellular activities (AAA-ATPase), which removes the ubiquitinated proteins from the membrane before transferring them to the proteasome for degradation (Ruggiano et al., 2014; Christianson and Carvalho, 2022; Koch and Yu, 2019; Smoyer and Jaspersen, 2019; Mannino and Lusk, 2022).

In the fission yeast *Schizosaccharomyces pombe*, several INM proteins have been studied for their roles in the organization and regulation of chromosome stability, including Ima1, Lem2, Man1, Bqt3, and Bqt4 (Chikashige et al., 2006; Chikashige et al., 2009; Hiraoka et al., 2011; Steglich et al., 2012; Gonzalez et al., 2012; Tange et al., 2016; Hirano et al., 2020b). In particular, we found that Bqt4 was degraded in the absence of Bqt3 (Chikashige et al., 2009). Bqt3 is a transmembrane protein composed of seven transmembrane domains (TMDs) along its entire length. Bqt4, which interacts with Bqt3, is predicted to be a tail-anchored protein. Immunoelectron microscopy analysis has shown that Bqt3 and Bqt4 localize at the INM (Chikashige et al., 2009). Bqt3 and Bqt4 are required for anchoring telomeres to the NE during vegetative growth and clustering telomeres to the spindle pole body through the meiosis-specific telomere-binding proteins Bqt1 and Bqt2 (Chikashige et al., 2006; Chikashige et al., 2009). Importantly, Bqt4 interacts with Lem2 (Hirano et al., 2018), and the depletion of Bqt4 is synthetic-lethal with the depletion of Lem2 (Tange et al., 2016; Kinugasa et al., 2019). Therefore, Bqt4 may play an important role in cell viability.

However, the non-telomeric vital roles of Bqt4 and the mechanisms underlying its degradation remain unknown. In this study, we identified the factors required for Bqt4 degradation through an INM-associated degradation mechanism. We also demonstrated that excess amounts of Bqt4 resulted in impaired nuclear morphology and growth defects, indicating the importance of regulated degradation of Bqt4 for membrane homeostasis.

## Results

### INM protein Bqt4 is degraded via the UPS

Because the UPS accounts for the selective degradation of many proteins in the nucleus (Enam et al., 2018; Franić et al., 2021), we first tested the requirement of the proteasome for Bqt4 degradation. We treated cells expressing N-terminally GFP-tagged Bqt4 (GFP-Bqt4) under its own promoter with bortezomib (BZ), a chemical proteasome inhibitor working in *S. pombe* (Takeda et al., 2011), and assessed the fluorescence levels of GFP-Bqt4 at 4 h after drug addition in the presence or absence of Bqt3. As a result of the treatment, GFP-Bqt4 fluorescence in the nuclei of both *bqt3^+^* and *bqt3*Δ cells was elevated (Fig. 1A). To confirm this result, we measured protein levels by western blotting, which consistently showed an increase in GFP-Bqt4 protein levels in these strains upon proteasomal inhibition by BZ (Fig. 1B). Notably, the GFP-Bqt4 signal was restored upon proteasome inhibition in the absence of Bqt3 and also elevated in the presence of Bqt3. This suggests that Bqt4 is constitutively turned over by the proteasome system and its association with Bqt3 antagonizes this process. We sought to confirm this phenomenon by introducing temperature-sensitive *mts2-1* and *mts3-1* proteasome mutations (Gordon and McGurk, 1993; Gordon et al., 1996). Using both fluorescence microscopy and western blotting, we found that GFP-Bqt4 levels were increased in *bqt3^+^*and *bqt3Δ* cells carrying these proteasome mutations at a restrictive temperature of 36 ^°^C, as compared to the corresponding wild-type strains (Fig. 1C, D; Fig. S1). These results indicate that Bqt4 degradation proceeds via the proteasome system.

**Fig. 1.**
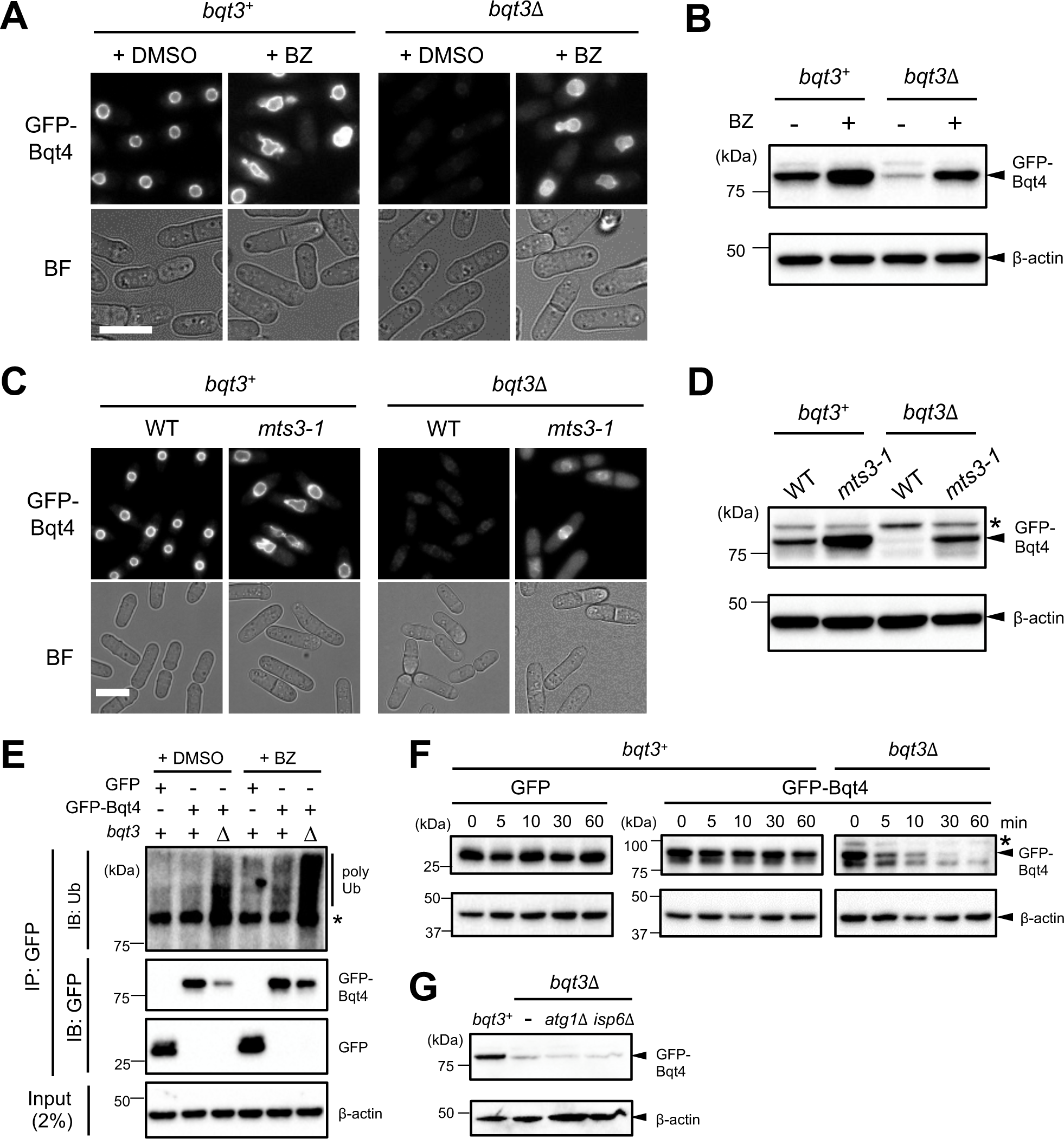
Bqt4 protein is degraded by the ubiquitin-proteasome system. (A, B) Cells expressing GFP-Bqt4 in the *bqt3*^+^ or *bqt3*Δ background were treated with vehicle (+DMSO) or the proteasome inhibitor bortezomib (+BZ) for 4 h and subjected to microscopic observation (A) or western blotting (B). (A) Fluorescence microscopy images of GFP-Bqt4 (upper panels); bright-field images (lower panels). Bar, 10 μm. (B) Protein amounts of GFP-Bqt4 and β-actin were detected by anti-GFP and anti-β-actin antibodies, respectively. Molecular weight markers are shown on the left. (C, D) Cells harboring *mts3-1* in the *bqt3*^+^ or *bqt3*Δ background were cultured at a nonpermissive temperature (36°C) for 4 h and subjected to microscopic observation (C) or western blotting (D). (C) Fluorescence microscopy images of GFP-Bqt4 (upper panels); bright-field images (lower panels). Bar, 10 μm. (D) Protein amounts of GFP-Bqt4 and β-actin were detected as in (B). The asterisk in (D) indicates non-specific bands. (E) Polyubiqutination of Bqt4 protein. Cells expressing GFP or GFP-Bqt4 in the *bqt3*^+^ or *bqt3*Δ background were treated with vehicle (+DMSO) or the proteasome inhibitor bortezomib (+BZ) for 4 h. After denaturing the proteins, GFP and GFP-Bqt4 were immunoprecipitated with anti-GFP antibody (IP). The precipitated proteins were detected by anti-ubiquitin or anti-GFP antibodies, respectively (IB). β-actin was used as a loading control. Molecular weight markers are shown on the left. (F) Bqt4 degradation rate in *bqt3*^+^ or *bqt3*Δ cells. The *bqt3*^+^ or *bqt3*Δ cells expressing GFP or GFP-Bqt4 were cultured in cycloheximide-containing YES medium for the time indicated on the top of the panels. The level of GFP and GFP-Bqt4 in the cells were detected by western blotting. β-actin was used as a loading control. Asterisk represents non-specific band. Molecular weight markers are shown on the left. (G) No Bqt4 degradation by the lysosome-autophagy system. Deletion of *atg1* (*atg1*Δ) or *isp6* (*isp6*Δ) was introduced into the *bqt3*Δ cells. Protein amounts of GFP-Bqt4 and β-actin in these cells were detected as in (B).Molecular weight markers are shown on the left.

Since proteins are ubiquitinated as a requirement for recognition by the proteasome, we examined whether Bqt4 was modified by polyubiquitination. We immunoprecipitated GFP-Bqt4 under denaturing conditions and detected associated ubiquitin by immunoblotting. Ubiquitinated forms of Bqt4 were detected in *bqt3*Δ cells and were enriched in both *bqt3^+^* and *bqt3*Δ cells when proteasomal activity was inhibited (Fig. 1E), suggesting that Bqt4 was targeted to the proteasome by polyubiquitin modification.

Bqt4 is degraded via the UPS in the absence of Bqt3, suggesting that Bqt3 plays a role in stabilizing Bqt4 by protecting Bqt4 from degradation. To assess whether Bqt3 stabilizes Bqt4, we determined the degradation kinetics of GFP-Bqt4 in the presence or absence of Bqt3 by cycloheximide chase assay followed by western blotting. The result showed that GFP-Bqt4 degraded faster in the absence of Bqt3 than in the presence of Bqt3 (Fig. 1F). This indicates that Bqt4 is protected from proteasome-dependent degradation by Bqt3. Thus, Bqt4 is maintained at the INM by associating with Bqt3, and is polyubiquitinated and degraded quickly once it dissociates from Bqt3.

Vacuolar autophagy is another pathway through which cells degrade proteins. Thus, we next examined whether Bqt4 degradation is regulated by the vacuolar autophagy pathway by analyzing GFP-Bqt4 levels in *bqt3*Δ cells lacking *isp6^+^* and *atg1^+^*genes. Isp6 is a serine protease necessary for vacuolar function, and Atg1 is a serine/threonine protein kinase that is the upstream initiator of the autophagy machinery. Deletion of these two genes has been shown to completely impair the vacuolar autophagy pathway (Kohda et al., 2007). We found that the deletion of *isp6* and *atg1* had no detectable effect on the quantity of GFP-Bqt4 in the absence of Bqt3, suggesting that Bqt4 degradation does not involve the vacuolar autophagy system (Fig. 1F). Collectively, we conclude that the Bqt4 protein is degraded by the UPS without the involvement of the vacuolar autophagy pathway.

### Bqt4 degradation occurs in the nucleus

Next, we determined whether Bqt4 was degraded in the nucleus by the nuclear proteasome. In *S. pombe*, proper targeting of proteasomes to the nucleus requires the tethering factor, Cut8 (Tatebe and Yanagida, 2000). In *cut8* mutants, proteasomes fail to localize to the nucleus and are mostly cytoplasmic (Tatebe and Yanagida, 2000). Because nuclear protein degradation depends on the localization of proteasomes in the nucleus (Takeda and Yanagida, 2005), we examined the levels of GFP-Bqt4 in *bqt3^+^*and *bqt3Δ* cells carrying the temperature-sensitive *cut8-563* mutation. We found that GFP-Bqt4 degradation was suppressed in *cut8-563* cells shifted to the nonpermissive temperature of 36°C (Fig. 2A, B). Taken together, our results strongly suggest that Bqt4 is degraded mainly by the proteasome within the nucleus.

**Fig. 2.**
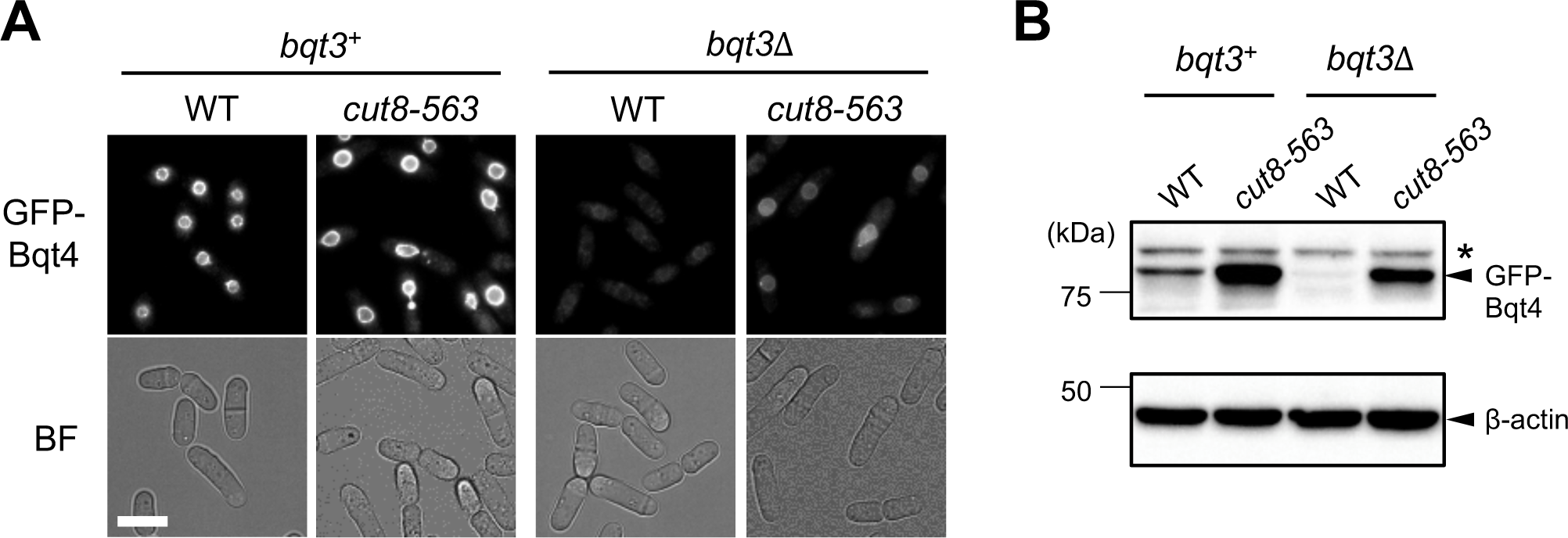
Bqt4 degradation occurs in the nucleus. Cells harboring *cut8-563 ts* mutation in the *bqt3*^+^ or *bqt3*Δ background were cultured at a nonpermissive temperature (36°C) for 4 h and subjected to microscopic observation (A) or western blotting (B). (A) Fluorescence images of GFP-Bqt4 (upper panels) and bright-field images (lower panels). Bar, 10 μm. (B) Protein levels of GFP-Bqt4 and β-actin were detected using anti-GFP and anti-β-actin antibodies, respectively. Asterisks indicate non-specific bands. The molecular weight markers are shown on the left.

### Bqt4 degradation involves the Doa10 E3 ligase complex

We next sought to identify the specific E3 ubiquitin ligase that directly recognizes Bqt4. As Bqt4 is a protein associated with the INM, we investigated whether membrane-localized Hrd1 and Doa10 E3 ligases participate in the Bqt4 degradation pathway. Deletion of *doa10^+^*partially restored GFP-Bqt4 levels in the absence of Bqt3, whereas deletion of *hrd1^+^* did not (Fig. 3A and B, comparing *hrd1*Δ to *doa10*Δ). These results are consistent with previous studies that showed that Doa10 partially localizes to the INM and is involved in the degradation of certain nuclear and INM substrates, whereas Hrd1 is found exclusively in the ER (Deng and Hochstrasser, 2006; Boban et al., 2014). Additionally, the double deletion of *doa10^+^* and *hrd1^+^* showed no obvious synthetic effect on GFP-Bqt4 restoration compared with the deletion of *doa10^+^* alone (Fig. 3A, B). We also measured the amount of GFP-Bqt4 in mutants lacking Ubc6 and Ubc7, which are E2 ubiquitin-conjugating enzymes associated with Doa10 (Swanson et al., 2001). We found that GFP-Bqt4 levels increased to some extent in *ubc6*1′, *ubc7*1′ single, and *ubc6*1′ *ubc7*1′ double mutants (Fig. 3C, D). These findings indicate that the Doa10 complex contributed to the degradation of Bqt4 in the INM.

**Fig. 3.**
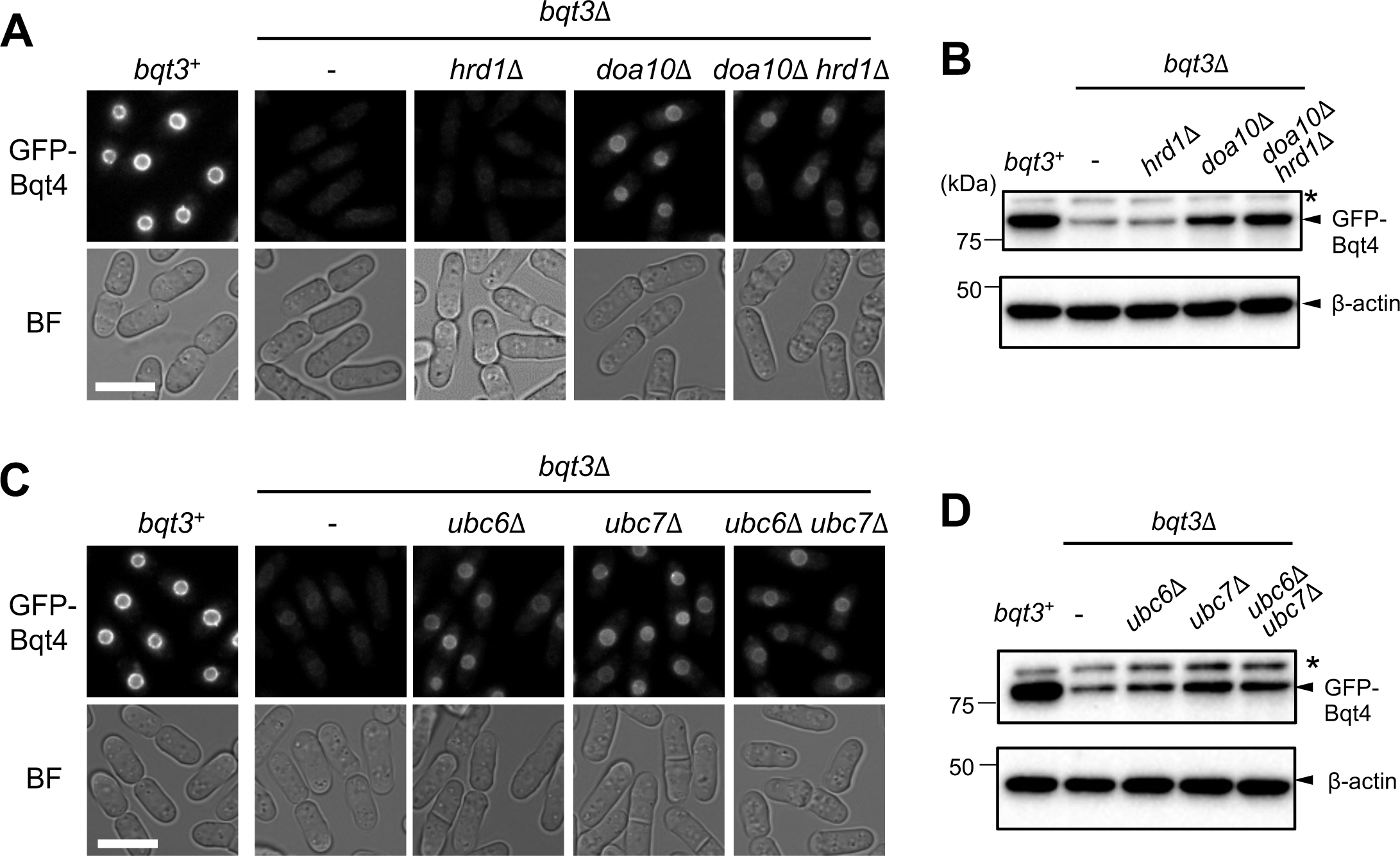
Bqt4 is a substrate of the Doa10 complex. (A, B) Cells harboring *hrd1* (*hrd1*Δ), *doa10* (*doa10*Δ) single deletion, or *hrd1* and *doa10* double deletion (*hrd1*Δ *doa10*Δ) in *bqt3*Δ background were cultured at 30°C for 16 h, and then subjected to microscopic observation (A) or western blotting (B). (A) Fluorescence images of GFP-Bqt4 (upper panels) and bright-field images (lower panels). Bar, 10 μm. (B) Protein levels of GFP-Bqt4 and β-actin were detected using anti-GFP and anti-β-actin antibodies, respectively. (C, D) Cells harboring *ubc6* (*ubc6*Δ), *ubc7* (*ubc7*Δ) single deletion, or *ubc6* and *ubc7* double deletion (*ubc6*Δ *ubc7*Δ) in *bqt3*Δ background were cultured at 30°C for 16 h, and then subjected to microscopic observation (C) or western blotting (D). (C) Fluorescence images of GFP-Bqt4 (upper panels) and bright-field images (lower panels). Bar, 10 μm. (D) Protein levels of GFP-Bqt4 and β-actin were detected using anti-GFP and anti-β-actin antibodies, respectively. Asterisks indicate non-specific bands. The molecular weight markers are shown on the left.

Although degradation of Bqt4 depended on Doa10, the level of GFP-Bqt4 was only partially restored in the *doa10*1′ mutant (Fig. 3A, B). Therefore, we attempted to identify other redundant E3 ligases involved in Bqt4 degradation. We constructed mutants of E3 ligases and their components that have been suggested to localize to or function in the ER and nucleus, namely *meu34*1′, *dsc1*1′, *ubr1*1′, *ubr11*1′, *san1*1′, *hul5*1′, *ptr1-1*, and *cut9-665* (PomBase (https://www.pombase.org), Matsuyama et al., 2006, Nielsen et al., 2014; Enam et al., 2018; Franić et al., 2021), and observed GFP-Bqt4 fluorescence intensity in these mutants. The mutants tested showed no detectable increase in fluorescence, except for the *hul5*1′ mutant (Fig. S2). The *hul5*1′ cells showed a slight increase in the fluorescence level of GFP-Bqt4, but showed no increase in GFP-Bqt4 level in western blotting (Fig. S3). Instead, a small degraded fragment was observed (Fig. S3), suggesting that the increase in the GFP signal in *hul5*1′ cells reflects this degraded product. Therefore, Doa10 is the only E3 ligase identified to date that ubiquitinates Bqt4.

### Bqt4 is an integral membrane protein

We have previously shown that Bqt4 localizes exclusively to the nucleoplasmic side of the INM, and this localization requires its C-terminal helix (Chikashige et al., 2009). However, this C-terminal helix was predicted to be a weak candidate for TMD (Fig. S4), we examined whether Bqt4 is an integral or peripheral membrane protein to understand how Bqt4 is removed from INM before degradation. Therefore, we performed membrane protein extraction experiments. In brief, the membrane fraction was isolated from the cell extract, treated under various conditions, and then separated into pellet and supernatant fractions by ultracentrifugation, followed by western blotting; a protein was expected to enter the pellet fraction if it remained in the lumen or associated with the membrane after the treatment. GFP-ADEL (Pidoux & Armstrong, 1992) and Ish1-GFP were used as controls for the luminal and integral membrane proteins, respectively. As shown in Fig. 4A, all these proteins were extracted from the membrane by treatment with Triton X-100 and 0.5 M NaCl. However, similar to Ish1, Bqt4 did not dissociate from the membrane by sodium carbonate (pH 11.5) or urea, which can strip peripheral membrane proteins. In contrast, GFP-ADEL was released from the membrane following sodium carbonate treatment, as expected for luminal proteins. Thus, Bqt4 is likely to behave as a tail-anchored protein integrally embedded in the INM in the presence of Bqt3.

**Fig. 4.**
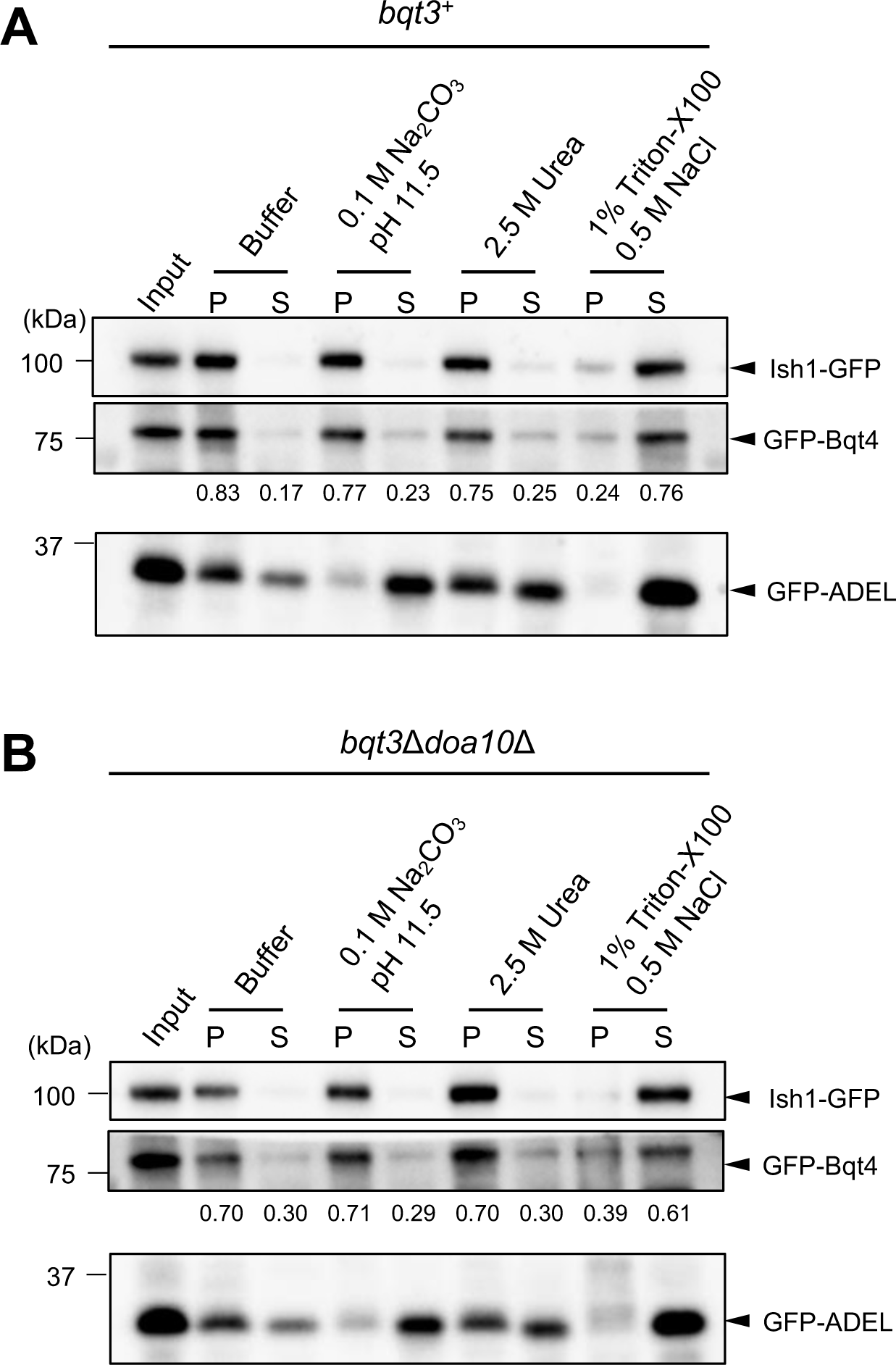
Bqt4 is an integral membrane protein. (A) Cells expressing GFP-Bqt4, Ish1-GFP, or GFP-ADEL were mixed and lysed by bead beating. After isolating membrane fraction, the membrane was treated under various conditions as shown on the top. The extracts were fractionated by ultracentrifugation into supernatant (S) and pellet (P), and then GFP-tagged proteins in each fraction were detected by western blotting. Molecular weight markers are shown on the left. Values at the middle panel represent the GFP-Bqt4 intensity ratio in supernatant and pellet fractions. (B) The *bqt3*Δ *doa10*Δ cells expressing GFP-Bqt4 were treated as described in A and then GFP-tagged proteins in each fraction were detected by western blotting. Molecular weight markers are shown on the left. Values at the middle panel represent the GFP-Bqt4 intensity ratio in supernatant and pellet fractions.

To further characterize the properties of TMD, we examined whether Bqt4 behaves as an integral membrane protein in the absence of Bqt3 using a membrane extraction assay. As most of Bqt4 protein is degraded in the *bqt3*Δ cells, we used the *bqt3*Δ*doa10*Δ cells to prevent the degradation. Bqt4 was extracted from the membrane by detergent, but not by sodium carbonate and urea (Figure 4B), as was the case in the *bqt3*^+^ condition, indicating that the TMD of Bqt4 was embedded in the INM independent of Bqt3.

### Degradation of Bqt4 requires the Cdc48 retrotranslocation complex

The ATPase Cdc48 has been shown to facilitate the degradation of ubiquitinated ERAD membrane substrates by extracting them from the ER membrane in *S. cerevisiae* cells (Ye et al., 2001; Jarosch et al., 2002; Rabinovich et al., 2002; Wolf and Stolz, 2012). Therefore, we investigated whether Cdc48 is necessary for Bqt4 degradation. To this end, we generated a *cdc48* mutant bearing mutations in both the D1 (P267L) and D2 (A550T) ATPase domains based on the temperature-sensitive *cdc48-6* allele in *S. cerevisiae* (Schuberth and Buchberger, 2005; Ruggiano et al., 2016) (Fig. 5A). The *cdc48* mutant (*cdc48-P267L/A550T*) exhibited cold-sensitive growth defects (Fig. 5B), suggesting that these mutations impaired Cdc48 activity in *S. pombe*. Inactivation of *cdc48-P267L/A550T* by shifting to a nonpermissive temperature of 16°C elevated the level of GFP-Bqt4 in *bqt3^+^* and *bqt3*1′ cells, as observed under fluorescence microscopy (Fig. 5C) and western blotting (Fig. 5D); however, it was degraded at 33°C (Fig. S5), indicating that Cdc48 was required for Bqt4 degradation.

**Fig. 5.**
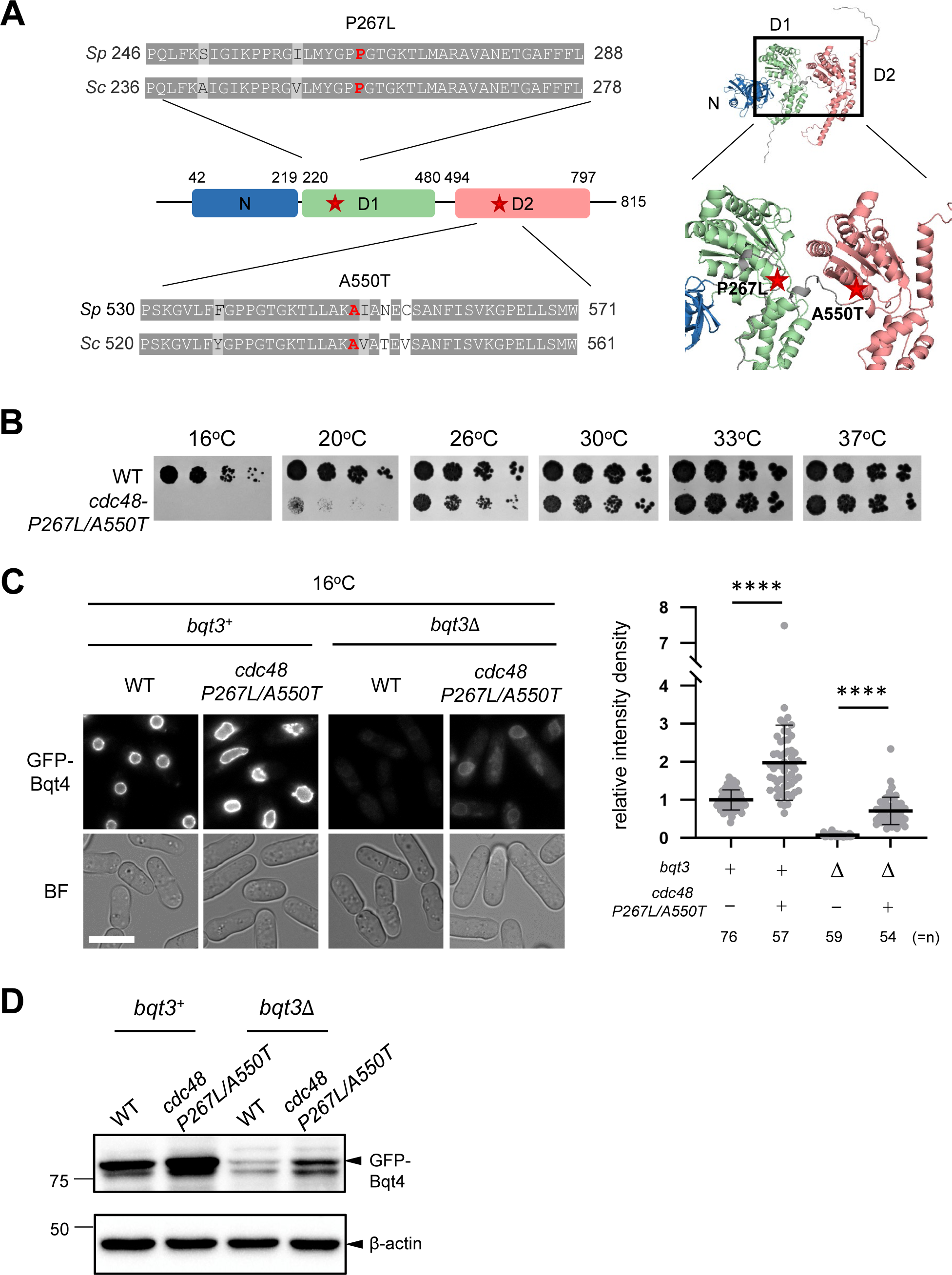
Removal of Bqt4 requires the Cdc48 retrotranslocation complex. (A) The Cdc48 mutant used in this study. Left panel: the amino acid sequence alignment around the D1 and D2 domains of *S. pombe* (*Sp*) and *S. cerevisiae* (*Sc*) Cdc48. Gray and light gray shades denote identical and similar amino acids, respectively (top and bottom). The amino acids indicated by red (P267 and A550; correspond to red stars in the schematic diagram in the middle) are both mutated in this study (P267L/A550T). Right panel: the mutation sites are shown in the predicted three-dimensional structure of Cdc48 by AlphaFold2 (https://alphafold.ebi.ac.uk/entry/Q9P3A7). (B) Temperature sensitivity of *cdc48-P267L/A550T* mutant. Five-fold serially diluted cells harboring *cdc48-P267L/A550T* mutant were spotted on the YES plate, and cultured at different temperatures as indicated at the top (C, D) Effect of hypomorphic mutation of *cdc48* on Bqt4 degradation. Cells harboring the *cdc48-P267L/A550T* mutant in the *bqt3*^+^ or *bqt3*Δ background were cultured at a nonpermissive temperature (16°C) for 15 h and subjected to microscopic observation (C) or western blotting (D). (C) Left panels: Fluorescence images of GFP-Bqt4 (upper panels); bright-field images (lower panels). Bar: 10 μm. Right panel: the fluorescence intensities in the nuclei were quantified and the relative values were plotted. The mean values are plotted with standard deviation on the right panel. The number of the cells analyzed is shown at the bottom. ∗∗∗∗*p* < 0.0001 via two-tailed unpaired student’s *t*-test. (D) Protein amounts of GFP-Bqt4 and β-actin were detected by anti-GFP and anti-β-actin antibodies, respectively. Molecular weight markers are shown on the left.

### The C-terminal TMD of Bqt4 acts as a degradation signal

We previously showed that Bqt4 degradation depends on its C-terminal TMD (residues 414-432) (Chikashige et al., 2009). To identify the minimal region of Bqt4 that is sufficient for its degradation, we analyzed truncated C-terminal fragments based on the structure of Bqt4 predicted by AlphaFold2 (Fig. 6A, Fig. S6). The Bqt4 fragment containing the helix domain and the adjacent intrinsically disordered sequence (Bqt4C^369-432^) reproduced the behavior of full-length Bqt4, as shown in Fig. 1A, that is, localization to the NE and responses to BZ, both in the presence and absence of Bqt3 (Fig. 6A, Bqt4C^369-432^). The helix domain alone (Bqt4C^394-432^) showed weaker localization to the NE and was somewhat diffused to the membrane compartments in the cytoplasm; nevertheless, this fragment was degraded in the absence of Bqt3, and its level increased with BZ treatment (Fig. 6A, Bqt4C^394-432^). The TMD alone (Bqt4C^414-432^) showed even weaker localization at the NE with diffusion to the cytoplasm, but showed the same responses, that is, degradation in the absence of Bqt3 and increased levels upon BZ treatment (Fig. 6A, Bqt4C^414-432^). These results indicate that the C-terminal fragment containing residues 369–432 was necessary and sufficient for the NE localization of Bqt4, and its truncation successively reduced NE localization. These results also indicate that the C-terminal TMD of Bqt4 is sufficient for its proteasome-mediated degradation.

**Fig. 6.**
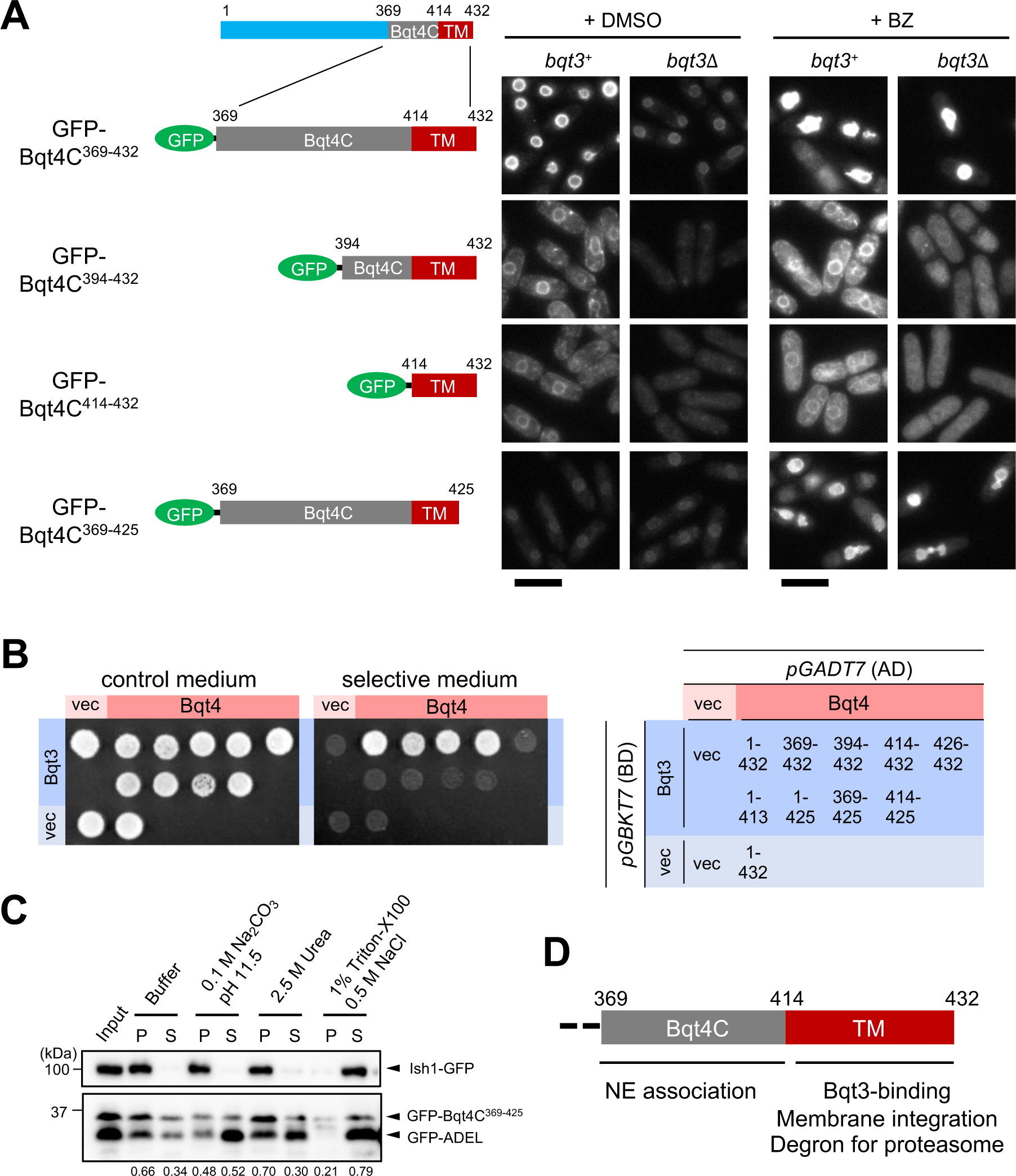
The C-terminal sequence of Bqt4 is a degron. (A) The C-terminal fragments of Bqt4 are shown as a cartoon on the left. These fragments were expressed as GFP-fused proteins in the *bqt3*^+^ or *bqt3*Δ cells. The cells were treated with vehicle (+DMSO) or the proteasome inhibitor bortezomib (+BZ) for 4 h and then observed by fluorescence microscopy. Bars: 10 μm. (B) Interaction between C-terminal truncated fragments of Bqt4 and full-length Bqt3 was examined by the yeast two-hybrid assay. AD, the transcriptional activation domain only (vec) or fused to GFP-Bqt4 fragments were expressed from the pGADT7 plasmid. BD, the DNA-binding domain only (vec) or fused to full-length Bqt3 were expressed from the pGBKT7 plasmid. The combination of AD- and BD-fused fragments are summarized at the right. The cells harboring both plasmids were cultured on control and selective medium. (C) Membrane association assay of GFP-Bqt4C^369-425^. The fragments were extracted from the membrane as described in Fig. 4A and detected by western blotting. S and P represent supernatant and pellet, respectively. Molecular weight markers are shown on the left. Values at the bottom represent the GFP-Bqt4 intensity ratio in supernatant and pellet fractions. (D) Schematic diagram of roles of the C-terminus of Bqt4.

To further narrow down the TMD of Bqt4 for proteasome-dependent degradation and Bqt3-dependent protection against degradation, we generated constructs containing a shorter TMD with the last seven residues truncated (Bqt4C^369-425^). The fragment Bqt4C^369-425^ lost its dependency on Bqt3 for degradation and exhibited similar behaviors in the presence or absence of Bqt3 (Fig. 6A, Bqt4C^369-425^): this fragment showed increased signal levels in the presence of BZ, indicating that it is also a substrate of proteasomal degradation in a Bqt3-independent manner. As the level of Bqt4C^369-425^ were not restored by *doa10* deletion, the fragment is likely degraded by a distinct pathway from Doa10 (Supplementary Fig. S7). To further characterize the reason why these fragments were degraded independent of Bqt3, we examined the interaction between these fragments and Bqt3 as well as the membrane association state of these fragments. The results of the yeast-two-hybrid (Y2H) assay showed that the fragments lacking the last seven residues did not bind to Bqt3 (Fig. 6B). In addition, Bqt4C^369-425^ was partially extracted by sodium carbonate (Fig. 6C), suggesting that the last seven residues assisted the membrane-embedding ability of the TMD. Thus, we concluded that the entire TMD (414–432) is the minimal region of the Bqt3-binding site and is a degron for proteasome-dependent degradation that is protected by Bqt3 (Fig. 6D).

### Excess Bqt4 causes deformation of the NE

We observed that inhibition of proteasomal degradation caused striking NE expansion and deformation, with irregular protrusions and invaginations, especially in *bqt3^+^* cells, in which Bqt4 levels were excessively high (Fig. 1A, C). We hypothesized that high levels of Bqt4 in these cells were responsible for this abnormal NE morphology. To test this possibility, we inhibited proteasomal degradation in the presence or absence of endogenous Bqt4 and visualized the NE using Ish1-GFP as an NE marker, and then nuclear deformation was assessed by measuring the roundness of the nucleus. Only when Bqt4 was present, a strong deformation of the NE was observed upon proteasomal inhibition (see *bqt4*^+^+BZ; Fig. 7A), suggesting that Bqt4 was an essential factor causing this phenomenon. To ascertain whether this abnormal phenotype was caused by an accumulation of Bqt4, we overexpressed GFP-Bqt4 under the *nmt1* promoter using YAM2 to control the expression level. YAM2, a chemical compound, suppresses *nmt1* promoter activity depending on its concentration (Nakamura et al., 2011). Overexpression of GFP-Bqt4 reproduced the nuclear-deformed morphology (Fig. 7B), and NE proliferation and deformation became more striking in cells that expressed higher levels of GFP-Bqt4 (Fig. 7C, D), indicating that the abnormal NE morphology depended on the amount of Bqt4. Taken together, these results support our hypothesis that Bqt4 is the only protein whose accumulation upon inhibition of degradation induces aberrant nuclear morphology. Finally, we investigated whether an excess of GFP-Bqt4 was toxic. We found that overproduction of GFP-Bqt4 caused a growth defect in both *bqt3^+^*and *bqt3Δ* strains compared to cells expressing endogenous levels of Bqt4 (Fig. 7E). Altogether, these observations suggest that Bqt4 plays an important role in regulating NE morphology, and a nuclear proteasomal degradation pathway controlling the quantity of Bqt4 is crucial for nuclear membrane homeostasis.

**Fig. 7.**
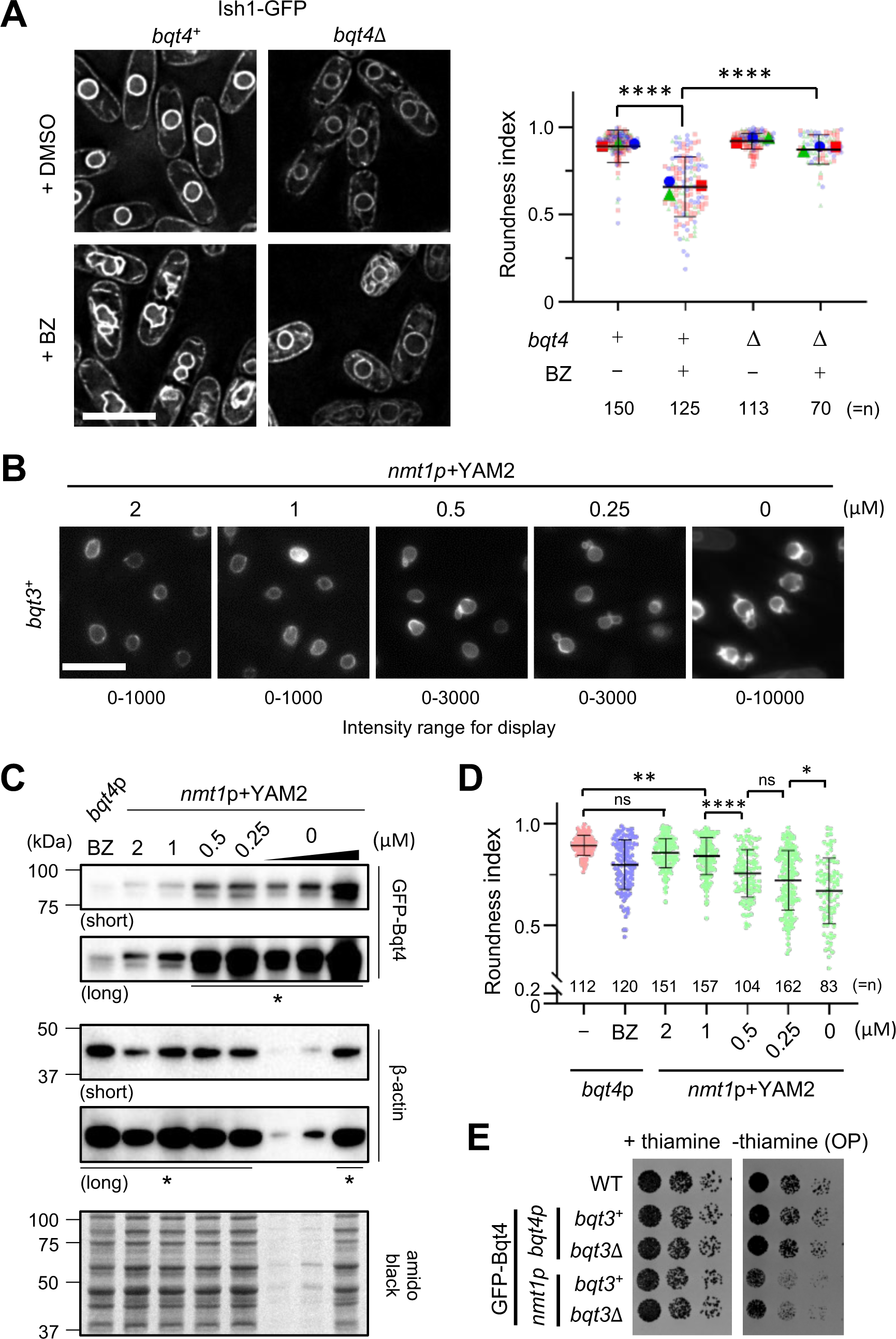
Excess Bqt4 causes nuclear envelope deformation. (A) Left panels: Fluorescence images of Ish1-GFP in *bqt4*^+^ or *bqt4*Δ cells. Ish1-GFP was used as a nuclear membrane marker (Asakawa et al., 2022). The cells were treated with vehicle (+DMSO) or the proteasome inhibitor bortezomib (+BZ) for 4 h. The images were deconvoluted using built-in SoftWoRx software with default settings. Bar: 10 μm. Right panel: The roundness index of the nuclei in these cells was measured from three independent experiments (blue, green, and red) and plotted as a superplot (Lord et al., 2020) (right panel). The roundness index of 1 indicates a perfect circle. The solid blue circle, green triangle, and red square represent the mean values in each experiment. Black horizontal lines represent the mean values for all data shown with standard deviations. *n* at the bottom right represents the total number of cells analyzed. ∗∗∗∗*p* < 0.0001 via two-tailed unpaired student’s *t*-test. (B-D) NE deformation due to Bqt4 overexpression. (B) Representative fluorescence images of GFP-Bqt4 in *bqt3*^+^ cells. GFP-Bqt4 was overexpressed under the control of the *nmt1* promoter in the presence or absence of YAM2 in the EMMG for 20 h. The YAM2 concentrations are shown in the images. To improve visibility, the contrast of the image was adjusted to the intensity range indicated at the bottom. Bar: 10 μm. (C) GFP-Bqt4 protein levels. Protein levels of GFP-Bqt4 shown in (B) were determined by western blotting. β-actin was used for loading control. Because the full expression of GFP-Bqt4 (0 μM YAM2) was strong, 1/10 (0.5 μg), 1/5 (1 μg), and 1 (5 μg) relative amounts of total proteins were electrophoresed. Short- and long-exposure images are shown in upper and lower panels, respectively. The bands indicated by asterisks in the bottom panel are saturated. The molecular weight markers are shown on the left. (D) Roundness index of the nuclei was mesured in BZ-treated (*bqt4*p, BZ; blue) and YAM2-treated (*nmt1*p+YAM2; green) *bqt3*^+^ cells overexpressing Bqt4. The roundness index was also measured in WT (*bqt4*p, -; red). The values were plotted with mean ± SD. *n* on the horizontal axis represents the number of cells analyzed. Significance was assessed using the Tukey’s test. ∗ *P* < 0.05; ∗∗∗∗ *P* < 0.0001; ns, not significant. (E) Growth defects by Bqt4 overproduction. GFP-Bqt4 was overexpressed in *bqt4* (*bqt4p*) or *nmt1* promoter (*nmt1p*) in *bqt3*^+^ or *bqt3*Δ cells. Five-fold serially diluted cells were spotted onto EMMG plates in the presence or absence of thiamine for four days. WT indicates wildtype.

## Discussion

In this study, we identified the molecular factors required for the degradation of the INM integral protein Bqt4 in *S. pombe* (Fig. 8). Bqt4 is ubiquitinated and degraded by the nuclear proteasome via an INMAD mechanism involving the Doa10 E3 ligase, which possibly recognizes the C-terminal TMD of Bqt4 as a degron. In addition, the AAA-ATPase Cdc48 is required for Bqt4 degradation. We also demonstrated that the accumulation of undegraded Bqt4 in the INM caused strong deformation of the NE and resulted in growth defects. Bqt4 escapes degradation by interacting with Bqt3 through its C-terminal TMD. Excess amounts of Bqt4 that were not bound to Bqt3 were subjected to INMAD-mediated degradation to avoid the accumulation of Bqt4 at the NE. These findings contribute to our understanding of the significance of INM protein degradation in INM homeostasis.

**Fig. 8.**
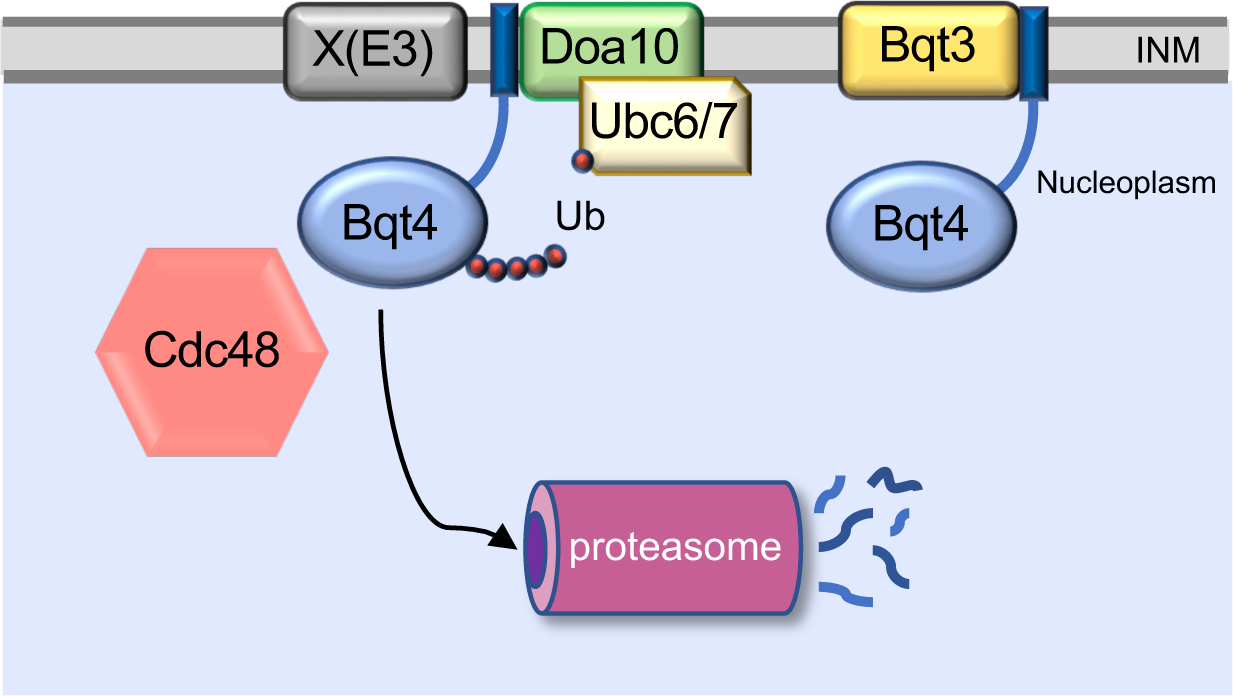
A model of INM protein degradation of Bqt4. Excess Bqt4, with the exposed TMD as a degradation signal, is recognized and ubiquitinated by INM-associated E3 ubiquitin ligase complexes, including Doa10 and additional unidentified E3 ligases. Ubiquitinated Bqt4 is then transferred to the nuclear proteasome for proteolysis. Degradation of Bqt4 requires the AAA-ATPase Cdc48 complex.

### Bqt4 is a native INM degradation substrate containing an intramembrane degron

Studies on mammalian cells have suggested that integral INM proteins are largely stable (Toyama et al., 2013; Buchwalter et al., 2019); however, it has been difficult to study their turnover. In the budding yeast *S. cerevisiae*, most of the INMAD substrates characterized thus far are not native INM proteins, but rather nuclear soluble proteins (Swanson et al., 2001; Ravid et al., 2006) and ER membrane proteins that are unassembled or mislocalized to the INM (Foresti et al., 2014; Khmelinskii et al., 2014; Natarajan et al., 2020). Thus, our study on Bqt4 provides an opportunity for a native INM substrate to obtain further insight into the mechanisms governing INM protein degradation. In addition to several previous studies that reported a handful of native INM substrates (Boban et al., 2014; Pantazopoulou et al., 2016; Smoyer et al., 2019; Koch et al., 2019), our results on the degradation of the INM protein Bqt4 demonstrate that INMAD is responsible for not only maintaining INM identity by removing foreign proteins but also for maintaining the proteostasis of INM integral components.

An important question is whether INMAD machinery specifically recognizes its substrates. Although a degradation signal has yet to be identified for integral INM substrates of INM-localized E3 ubiquitin ligases, such as Doa10 and the Asi complex, we identified that Bqt4 has an intramembrane degron. This suggests that INMAD may not simply detect and degrade misfolded or mislocalized proteins in the INM, but instead specifically recognizes a substrate to regulate its protein levels.

### An INM protein degradation pathway in *S. pombe*

In *S. cerevisiae*, the Doa10 E3 ubiquitin ligase cooperates with the Asi complex in INMAD (Koch and Yu, 2019; Smoyer and Jaspersen, 2019; Mannino and Lusk, 2022). As homologs of Asi proteins are absent in *S. pombe* as in other species, INMAD mechanisms may be diverse among organisms. However, increasing evidence has shown that the degradation of INM proteins in plants and mammals also employs the UPS and Cdc48/p97 (Tsai et al., 2016; Huang et al., 2020; Krshnan et al., 2022), suggesting a general mechanism for INM protein degradation.

In *S. cerevisiae*, Doa10 E3 ligase can passively diffuse from the ER to the INM and functions in both ERAD and INMAD (Deng and Hochstrasser, 2006). In particular, Doa10 has been implicated in mediating the degradation of the INM protein Asi2, which is a component of INMAD (Boban et al., 2014). In *S. pombe*, Bqt4 degradation also depends on Doa10, indicating that Bqt4 is subject to regulation by INMAD. In addition, our data indicate that *S. pombe* might possess additional unidentified redundant E3 ligases cooperating with Doa10 at the INM. Because Bqt4 is supposed to be recognized by these E3 ligases at its transmembrane degron, it is expected that the unidentified E3 ligases also localize to the INM (“X(E3)” in Fig. 8). Similar situations in which unidentified E3 ligases degrade an INM substrate have been reported in *S. cerevisiae* (Pantazopoulou et al. 2016). Similarly, INM-localized E3 ligases have yet to be identified in higher eukaryotes. We speculate that the uncharacterized E3 ligases functioning at the INM in *S. pombe* might operate with E2 ubiquitin-conjugating enzymes other than Ubc6 and Ubc7, because degradation of Bqt4 exhibits limited dependence on these enzymes. Thus, it would be of great interest to identify and characterize other membrane-associated E3 ligases and E2 conjugating enzymes that mediate protein degradation at the INM.

The Cdc48 complex plays a central role in the ERAD degradation of ER integral membrane proteins by exerting a pulling force on the ubiquitinated membrane protein via its ATPase activity (Braun, 2002; Rabinovich et al., 2002; Wolf and Stolz, 2012; Bodnar and Rapoport, 2017). Cdc48 has also been implicated in the degradation of INM proteins Asi1 and Msp3 in *S. cerevisiae* (Pantazopoulou et al. 2016; Koch et al., 2019). In *S. pombe*, Cdc48 mutations at its ATPase active center impair Bqt4 degradation (Fig. 5), indicating that ERAD/INMAD degradation involves the Cdc48 complex; however, it remains unclear whether Cdc48 extracts Bqt4 from the INM.

### The biological significance of Bqt4 degradation

The INM protein Bqt4 has been suggested to have non-telomeric functions that are currently unknown but are vital for cell growth, as the double deletion of Bqt4 and Lem2 results in synthetic lethality (Tange et al., 2016; Hirano et al., 2018; Kinugasa et al., 2019; Hirano et al., 2023b). In cells lacking both Bqt4 and Lem2, nuclear envelope rupture occurs in association with leakage of nuclear proteins into the cytoplasm, suggesting that Bqt4, in cooperation with Lem2, plays an essential role in maintaining nuclear membrane integrity (Kinugasa et al., 2019). The nuclear membrane rupture phenotype can be rescued by the increased expression of Elo2, a fatty acid elongase that catalyzes the synthesis of very-long-chain fatty acids (Kinugasa et al., 2019), and Tlc4, a ceramide synthase homolog which is localized in the Golgi (Hirano et al., 2023b). In addition, the depletion of Bqt4 and Lem2 leads to a reduction in the quantity of t20:0/24:0 phytoceramide, the most abundant ceramide species in *S. pombe*, and this reduction can be prevented by the overexpression of these proteins (Kinugasa et al., 2019; Hirano et al., 2023b). These findings strongly suggest that Bqt4 is involved in nuclear membrane lipid metabolism. Thus, we propose that the abnormal accumulation of Bqt4 at the INM may result in excessive lipid synthesis, ultimately leading to nuclear membrane expansion and deformation. One possibility is that Bqt4 promotes the synthesis of phospholipids, a major component of the nuclear membrane, as Bqt4 binds to an enzyme involved in lipid synthesis (Hirano et al, 2023a). Several lines of evidence suggest that increased phospholipid synthesis from phosphatidic acid induces striking NE membrane deformation (Santos-Rosa et al., 2005; Barbosa et al., 2015). The exact mechanisms leading to the deformed nuclear morphology upon the accumulation of Bqt4 remain to be elucidated. Further investigations will advance our understanding of the role of INM protein degradation in the maintenance of nuclear membrane homeostasis.

## Materials and methods

### Yeast strains and culture media

All *S. pombe* strains used in this study are listed in Table S1. Unless otherwise specified, all *S. pombe* strains were cultured in yeast extract with supplements (YES) medium. Malt extract medium (ME) was used to induce sporulation. G418 disulfate (Nacalai Tesque, Japan), hygromycin B (Wako, Japan), nourseothricin sulfate (Werner BioAgents, Germany), and blasticidin S (Wako, Japan) were used at concentrations of 100, 200, 100, and 100 μg/ml, respectively, to select strains bearing the respective selection marker.

### Construction of plasmids and strains

All plasmids were constructed using the NEBuilder system (New England BioLabs, MA, USA) and enzymatic digestion and ligation (TaKaRa, Japan) systems, according to the manufacturer’s instructions. Gene disruption and GFP tagging were performed using the direct chromosomal integration method (Wach, 1996; Bähler et al., 1998). The integration cassettes were amplified using a two-step PCR. In the first PCR, fragments (∼500 bp genomic regions upstream and downstream of the gene of interest) were amplified from the wild-type *S. pombe* genome using KOD One DNA polymerase (TaKaRa Bio Inc., Japan). The first PCR products were used as primers in the second PCR step to amplify a cassette for integration, which contained a selection marker. For gene disruption, pFA6-kanMX6 (Bähler et al., 1998), pCR2.1-hph, pCR2.1-bsd, and pCR2.1-nat (Sato et al., 2005) were used to generate the integration cassettes. For GFP tagging, pFA6-GFP(S65T)-kanMX6 (Bähler et al., 1998) was used to generate the GFP-containing integration cassettes. The cassettes were then transformed into *S. pombe* cells and the desired integrant strains were selected on YES plates containing the appropriate selection agent. Correct disruption and integration were confirmed by genomic PCR at the 5′ and 3′ ends using KOD FX Neo (TaKaRa Bio Inc., Japan). We confirmed that N-terminally GFP-tagged Bqt4 is functional, as it suppressed the synthetic lethality of *bqt4* and *lem2* double deletion (Tange et al., 2016) and anchored telomeres to the NE (Chikashige et al., 2009).

### Fluorescence microscopy

All fluorescence microscopy images were obtained from live cells. Fluorescence microscopy data of intracellular GFP fusion proteins were collected using a DeltaVision microscope system (GE Healthcare Inc., USA) equipped with a pco.edge 4.2 sCMOS camera (PCO, Germany) and a ×60 PlanApo N OSC oil-immersion objective lens (numerical aperture NA=1.4, Olympus, Japan). Cells were precultured from fresh colonies and cultured in YES medium at 30°C to reach the logarithmic growth phase (5–10×10^6^ cells/ml). The cells were then attached to glass-bottomed culture dishes coated with soybean lectin (Sigma-Aldrich, USA), covered with Edinburgh minimal medium with glutamate (EMMG) medium, and observed at specified temperatures.

### Image quantification and processing

An identical fluorescence intensity range was applied to all images presented in each panel using the Fiji software (Schindelin et al., 2012).

To quantify the fluorescence intensity in the nucleus, we used the Fiji software. First, we generated the sum projection image from the raw z-stack images and then subtracted the background signal from the entire image by measuring the mean intensity of a region where no cells were present. Second, the nuclear region was selected manually or semi-automatically. For semi-automatic selection, individual nuclei were selected (original image), and the selected image was blurred using Gaussian blur (blurred image). An appropriate sigma value was applied depending on the image contrast. An edge-enhanced image was generated by subtracting the blurred image from the original image. A threshold was applied to the nuclear region using the appropriate minimum and maximum intensity ranges. Third, the threshold region was applied to the original image and the total fluorescence intensity of the region was quantified. The nuclear intensities obtained from *bqt3*^+^ cells were averaged and the values were used to calculate the relative fluorescence intensity. The relative fluorescence intensities were plotted, and the results are presented as the mean ± standard deviation.

For the roundness index measurement, after selecting individual nuclei, a threshold was applied to the nuclear membrane signal using the appropriate fluorescence intensity ranges. The nuclear region was filled in, and the roundness index was measured using the built-in function in Fiji.

### Western blotting

Cells were precultured in YES medium at 30°C overnight, inoculated in fresh YES, and then cultured until the cell concentration reached mid-log phase (5–10×10^6^ cells/ml). Approximately 5×10^7^ cells were harvested by centrifugation at 4°C, and the cell pellet was washed with 1 ml ice-cold water containing 1 mM phenylmethylsulfonyl fluoride (PMSF). The cells were lysed in 150 μl of ice-cold 1.85 M NaOH containing 7.5% β-mercaptoethanol (β-ME) on ice for 15 min, followed by the addition of 150 μl of ice-cold 55% (v/v) trichloroacetic acid (TCA) to precipitate the proteins, incubation on ice for 15 min, and centrifugation (10 min, 14,000 ×g, 4°C). The protein pellet was then washed twice with ice-cold acetone and dissolved in sample solubilization buffer [50 mM Tris-HCl (pH 8.0), 1% SDS, 1× protease inhibitor cocktail (P8215, Sigma-Aldrich, USA), 1 mM PMSF, and 1 mM bortezomib]. Total protein concentrations were determined using the BCA assay (Thermo Fisher Scientific, USA) according to the manufacturer’s protocol. The samples were further diluted in 2× Laemmli SDS sample buffer to a concentration of 1 μg/μl. After the addition of dithiothreitol to a concentration of 50 mM, the samples were heated at 95°C for 10 min before immunoblotting. Identical amounts of protein (10 μg) were separated by SDS–PAGE and transferred to a polyvinylidene difluoride membrane. The membranes were blocked for 1 h at room temperature (approximately 26–28°C) in blocking solution [PBST (10 mM Na_2_HPO_4_, 137 mM NaCl, 2.7 mM KCl, 1.76 mM KH_2_PO_4_, and 0.1% Tween 20) containing 5% skim milk (Nacalai Tesque, Japan)]. To detect the GFP fusion proteins and β-actin as a loading control, the membranes were probed with anti-GFP rabbit polyclonal antibody (Rockland, USA; 1:2,000 dilution) or anti-β-actin mouse monoclonal antibody (Abcam, UK; 1:2,000 dilution) in blocking solution overnight at 4°C. After three rounds of washing with PBST, the membranes were incubated with appropriate horseradish peroxidase-conjugated secondary antibodies (1:2,000 dilution, Cytiva, USA) in a blocking solution at room temperature for 2 h. After three rounds of washing with PBST, the immunoreactive bands were detected using a chemiluminescence system (ImmunoStar LD; Wako, Japan) on a ChemiDoc imaging system (Bio-Rad, USA).

### Proteasome inhibition

Cells were grown in YES medium until early mid-log phase (2–5×10^6^ cells/ml), as described above. The culture was then divided into two groups, and the proteasome inhibitor bortezomib (LC Laboratories, USA) dissolved in dimethyl sulfoxide (DMSO) was added to one culture to a final concentration of 1 mM; the same amount of DMSO was added to the other culture. The cells were incubated at 26°C for 4 h before harvesting for fluorescence microscopy and immunoblotting.

### Temperature-sensitive mutant experiments

For the temperature-sensitive mutant experiments, cells harboring *ts* or *cs* mutations were grown in YES medium until the early mid-log phase (2–5×10^6^ cells/ml) at 26°C and 33°C, respectively. The culture temperature was shifted up to 36°C or down to 16°C, and then the cells were incubated for 4 h and 15 h, respectively, before harvesting.

### Ubiquitination detection assay

The ubiquitination detection assay was performed as described previously (Boban et al., 2014) with modifications. Cells were grown in YES medium at 30°C until the mid-log phase (5–10×10^6^ cells/ml), and approximately 5×10^8^ cells were harvested by centrifugation at 4°C. Where indicated, the proteasome inhibitor bortezomib was added at a final concentration of 1 mM for 4 h before the cells were collected. The same volume of DMSO was added to the control samples. The cells were washed with cold water containing 1 mM PMSF and lysed in 250 μl of 1.85 N NaOH solution containing 7.5% β-ME for 15 min on ice. Subsequently, the proteins were precipitated by adding 250 μl of 55% (v/v) TCA, followed by incubation for 15 min on ice. The proteins were pelleted by centrifugation (10 min, 14,000 ×g, 4°C) and washed with 500 μl of 50 mM Tris before being dissolved in 200 μl denaturation solution [25 mM Tris-HCl pH 7.4, 2% SDS, 2 mM ethylenediaminetetraacetic acid (EDTA), 1× protease inhibitor cocktail (P8215, Sigma-Aldrich, USA), 1 mM PMSF, 1 mM bortezomib, 10 μM PR-619 (SML0430, Sigma-Aldrich, USA), and 50 mM 2-chloroacetamide (032-09762, Fuji film Wako, Japan)]. Total protein concentrations were determined using the BCA assay (Thermo Fisher Scientific, USA) according to the manufacturer’s protocol. The samples were incubated at 65°C for 10 min and clarified by centrifugation (5 min, 15,000 ×g, room temperature). Equal amounts of protein lysates were diluted five-fold with cold IP dilution buffer (50 mM Tris-HCl pH 7.5, 1.2% Triton X-100, 2 mM EDTA, 100 mM NaCl, 1 × protease inhibitor cocktail, 1 mM PMSF, 1 mM bortezomib, 10 μM PR-619, and 50 mM 2-chloroacetamide), incubated with gentle rotation at 4°C for 1 h, and cleared by centrifugation at 14,000 ×g for 10 min. A portion (20 μl) of the sample (2% input) was mixed with 2× Laemmli SDS sample buffer (20 μl) and frozen until analysis. The remaining sample was mixed with antibody-coupled beads prepared from 2 μg anti-GFP rabbit polyclonal antibody (Rockland) and 50 μl protein-A-coupled dynabeads (Life Technologies, USA), according to the manufacturer’s instructions. The samples were immunoprecipitated at 4°C for 3 h with gentle rotation, and then pelleted using a magnet. The pellets were then washed as follows: twice with washing solution A (50 mM Tris-HCl pH 7.4, 2 mM EDTA, 100 mM NaCl, 1% Triton X-100, 0.4% SDS), once with washing solution B (50 mM Tris-HCl pH 7.4, 2 mM EDTA, 350 mM NaCl, 1% Triton X-100) including a 10-min incubation at 4°C with gentle rotation, and finally once with washing solution A. To elute the immunoprecipitated protein, 50 μl of 1× Laemmli SDS sample buffer was added to the pellet, and the mixture was incubated for 10 min at 65°C. Ten microliters of each sample were separated on SDS-PAGE gels (8%). For the analysis of immunoprecipitated GFP-Bqt4, the eluate was diluted 1:20 with 1× Laemmli SDS sample buffer. Immunoblotting was performed using an anti-GFP mouse monoclonal antibody (JL8; 1:1000; TaKaRa Bio Inc.) and an anti-ubiquitin mouse monoclonal antibody (P4D1, 1:1000; Santa Cruz Biotechnology, USA). Protein levels in the input samples were assessed using an anti-β-actin antibody (1:2000; Abcam).

### Cycloheximide chase assay

The *bqt3*^+^ or *bqt3*Δ cells expressing GFP or GFP-Bqt4 were grown in YES medium to the mid-log phase (approximately 5×10^6^ cells/ml). After adding final 200 μg/ml cycloheximide (C7698, Sigma-Aldrich), the cells were cultured for 0, 5, 10, 30, and 60 min for protein degradation. Immediately after culturing, stop solution (2 mM PMSF + 5× protease inhibitor cocktail) was added to the cells, and the cells were placed on ice for 5 min. The cells were harvested by centrifugation (8,000×g for 3 min) and subjected to western blotting as described above.

### Membrane association assay

The membrane association assay was performed according to a previously published protocol (Habeck et al., 2015) with slight modifications. *S. pombe* cells expressing GFP-Bqt4 and control proteins (Ish1-GFP and GFP-ADEL) were grown to the mid-log phase and mixed at an appropriate ratio before cell harvesting by centrifugation at 4°C. The harvested cells (∼5×10^8^ cells) were washed with cold water containing 1 mM PMSF and resuspended in 1 ml cold fractionation buffer (FB; 0.7 M sorbitol, 50 mM Tris-HCl pH 7.4, 150 mM NaCl, 1× protease inhibitor cocktail, 1mM PMSF, 1 mM bortezomib). Resuspended cells were lysed using a bead shocker (2,700 rpm for 10 cycles of 60 s on and 60 s off; Yasui Kikai, Co., Japan), followed by washing the beads three times with 200 μl FB. The cell extracts were combined, and cell debris and intact nuclei were removed by centrifugation (2000 ×g for 5 min at 4°C). Membrane fraction was isolated by ultracentrifugation at 100,000 ×g for 1 h at 4°C. The membrane was thoroughly resuspended in FB and divided into four separate aliquots that were subjected to one of the following treatments: (1) FB alone; (2) 0.1 M Na_2_CO_3_, pH 11*.5*; (3) 2.5 M urea; or (4) 1% Triton X-100 and 0.5 M NaCl. After incubation for 1 h on ice with occasional mixing, the samples were separated into pellet (P) and supernatant (S) fractions by ultracentrifugation at 100,000 ×g for 45 min at 4°C. The pellets were washed once with appropriately supplemented FB by centrifugation (100,000 ×g for 45 min at 4°C) before being resuspended in FB. Proteins were then precipitated from the pellet and supernatant fractions by adding TCA to a final concentration of 20% (v/v). After 30 min of incubation on ice, the proteins were pelleted by centrifugation (16,000 ×g for 10 min at 4°C), washed twice with ice-cold acetone, and dissolved in 1× Laemmli SDS sample buffer. Samples were incubated for 10 min at 95°C before immunoblotting.

### Yeast two-hybrid assay

The yeast two-hybrid assay was performed as described in our previous paper, with slight modifications (Asakawa et al., 2022). GFP-Bqt4 fragments and Bqt3 were expressed from pGADT7 and pGBKT7 plasmids in the budding yeast strain (AH109). To examine protein interactions, suspensions of the transformants were spotted onto SD plates with dropout supplements: Leu/-Trp (cat #630417) or -Ade/-His/-Leu/-Trp (cat #630428) (Takara Bio). The strains used in this assay are listed in Table S1.

### Spot assay

Cells expressing GFP-Bqt4 under the control (*bqt4p*) or nmt1 promoter (*nmt1p*) were pre-cultured in YES medium overnight. Five-fold dilution series of the cells (starting concentration was 8.0×10^5^ /mL) were spotted on the EMMG medium plates supplemented with or without 10 μM thiamine, and then the plates were incubated at 30°C for 4 days.

For the temperature-sensitivity growth assay, wild-type and Cdc48 mutant cells were spotted in a five-fold dilution series and incubated at 16, 20, 26, 30, 33, and 37°C for 3–6 days.

## Acknowledgements

We thank Chizuru Ohtsuki and Yoshino Kubota for their technical assistance; We also thank to Professor Mitsuhiro Yanagida (OIST) and the National BioResource Project Yeast in Japan for providing the *S. pombe* strains.

## Author contributions

Conceptualization: T.K.L., Y. Hirano, T.H., Y. Hiraoka; Methodology: T.K.L.,Y. Hirano, H.A., K.O., T.H., Y. Hiraoka; Validation: T.K.L, Y. Hirano, K.O., T.H., Y. Hiraoka; Formal analysis: T.K.L, Y. Hirano; Investigation: T.K.L, Y. Hirano, H.A.; Resources: K.O, T.F, Y. Hiraoka; Data curation: T.K.L., Y. Hirano, H.A., K.O., T.F., T.H., Y. Hiraoka; Writing - original draft: T.K.L, Y. Hirano, T.H., Y. Hiraoka; Writing - review & editing: T.K.L., Y. Hirano, H.A., K.O., T.F., T.H., Y. Hiraoka; Visualization: T.K.L., Y. Hirano; Supervision: T.H., Y. Hiraoka; Project administration: Y. Hiraoka; Funding acquisition: Y. Hirano, K.O., T.F., T.H., Y. Hiraoka.

## Data availability

Further information and requests for resources and reagents should be directed to Yasushi Hiraoka

---

**Supplementary Table 1.** Yeast strains used in this study.

| Strain | Genotype | Source | Figure |
| --- | --- | --- | --- |
| TL204 | *h<sup>-</sup> bqt4Δ::hph lys1<sup>+</sup>::bqt4p-GFP-bqt4* | This study | 1A-G, 2A, B, 3A-D, 4A, 5C, D 7C-E, S1, S3A, B, S5 |
| TL205 | *h<sup>-</sup> bqt4Δ::hph lys1<sup>+</sup>::bqt4p-GFP-bqt4 bqt3Δ::NAT* | This study | 1A-G, 2A, B, 3A-D, 5C, D, 7E, S1, S2, S3A, B, S5 |
| TL288 | *h<sup>-</sup> bqt4Δ::hph lys1<sup>+</sup>::bqt4p-GFP-bqt4 mts3-1* | This study | 1C, D |
| TL289 | *h<sup>-</sup> bqt4Δ::hph bqt3Δ::NAT lys1<sup>+</sup>::bqt4p-GFP-bqt4 mts3-1* | This study | 1C, D |
| H1N95 | *h<sup>-</sup> bqt4Δ::hph lys1<sup>+</sup>::bqt4p-GFP* | This study | 1E |
| H1N2114 | *h<sup>-</sup> bqt4Δ::kan<sup>r</sup> bqt3Δ::NAT aur1<sup>r</sup>::mCherry-bqt3 lys1<sup>+</sup>::bqt4p-GFP* | This study | 1F |
| TL233 | *h<sup>-</sup> bqt4Δ::hph bqt3Δ::NAT lys1<sup>+</sup>::bqt4p-GFP-bqt4 isp6Δ::bsd* | This study | 1G |
| TL234 | *h<sup>-</sup> bqt4Δ::hph bqt3Δ::NAT lys1<sup>+</sup>::bqt4p-GFP-bqt4 atg1Δ::kan<sup>r</sup>* | This study | 1G |
| TL290 | *h<sup>-</sup> bqt4Δ::hph lys1<sup>+</sup>::bqt4p-GFP-bqt4 cut8-563* | This study | 2A, B |
| TL291 | *h<sup>-</sup> bqt4Δ::hph bqt3Δ::NAT lys1<sup>+</sup>::bqt4p-GFP-bqt4 cut8-563* | This study | 2A, B |
| TL215 | *h<sup>-</sup> bqt4Δ::hph bqt3Δ::NAT lys1<sup>+</sup>::bqt4p-GFP-bqt4 doa10Δ::bsd* | This study | 3A, B, 4B, S2, S3A, B |
| TL219 | *h<sup>-</sup> bqt4Δ::hph bqt3Δ::NAT lys1<sup>+</sup>::bqt4p-GFP-bqt4 hrd1Δ::kan<sup>r</sup>* | This study | 3A, B, S2 |
| TL236 | *h<sup>-</sup> bqt4Δ::hph bqt3Δ::NAT lys1<sup>+</sup>::bqt4p-GFP-bqt4 doa10Δ::bsd hrd1Δ::kan<sup>r</sup>* | This study | 3A, B |
| TL220 | *h<sup>-</sup> bqt4Δ::hph bqt3Δ::NAT lys1<sup>+</sup>::bqt4p-GFP-bqt4 ubc7Δ::kan<sup>r</sup>* | This study | 3C, D |
| TL221 | *h<sup>-</sup> bqt4Δ::hph bqt3Δ::NAT lys1<sup>+</sup>::bqt4p-GFP-bqt4 ubc6Δ::kan<sup>r</sup>* | This study | 3C, D |
| TL266 | *h<sup>-</sup> bqt4Δ::hph bqt3Δ::NAT lys1<sup>+</sup>::bqt4p-GFP-bqt4 ubc6Δ::kan<sup>r</sup> ubc7Δ::bsd* | This study | 3C, D |
| HA1382-3D | *h<sup>-</sup> lys1-131 ura4-D18 leu1-32 bqt3Δ::LEU2 aur1<sup>r</sup>::bqt3p-GFP-bqt3* | This study | 4A |
| H1N429 | *h<sup>-</sup> lys1<sup>+</sup>::ish1-GFP* | This study | 4A, B, 6C, 7A |
| H1N789 | *h<sup>-</sup> lys1<sup>+</sup>::nmt41p-SS-GFP-ADEL* | This study | 4A, B, 6C |
| h-972 | *h<sup>-</sup>* | Laboratory stock | 5B, 7E |
| TL311 | *h<sup>-</sup> lys1-131 cdc48Δ::kan<sup>r</sup> aur1<sup>r</sup>::cdc48p-cdc48-P267L/A550T* | This study | 5B |
| TL306 | *h<sup>-</sup> bqt4Δ::hph lys1<sup>+</sup>::bqt4p-GFP-bqt4 cdc48Δ::kan<sup>r</sup> aur1<sup>r</sup>::cdc48p-cdc48-P267L/A550T* | This study | 5C, D, S5 |
| TL307 | *h<sup>-</sup> bqt4Δ::hph lys1<sup>+</sup>::bqt4p-GFP-bqt4 bqt3Δ::NAT cdc48Δ::kan<sup>r</sup> aur1<sup>r</sup>::cdc48p-cdc48-P267L/A550T* | This study | 5C, D, S5 |
| TL242 | *h<sup>-</sup> bqt4Δ::hph lys1<sup>+</sup>::bqt4p-GFP-bqt4C<sup>369-432</sup>* | This study | 6A |
| TL243 | *h<sup>-</sup> bqt4Δ::hph bqt3Δ::NAT lys1<sup>+</sup>::bqt4p-GFP-bqt4C<sup>369-432</sup>* | This study | 6A |
| TL246 | *h<sup>-</sup> bqt4Δ::hph lys1<sup>+</sup>::bqt4p-GFP-bqt4C<sup>394-432</sup>* | This study | 6A |
| TL247 | *h<sup>-</sup> bqt4Δ::hph bqt3Δ::NAT lys1<sup>+</sup>::bqt4p-GFP-bqt4C<sup>394-432</sup>* | This study | 6A |
| TL248 | *h<sup>-</sup> bqt4Δ::hph lys1<sup>+</sup>::bqt4p-GFP-bqt4C<sup>414-432</sup>* | This study | 6A |
| TL249 | *h<sup>-</sup> bqt4Δ::hph bqt3Δ::NAT lys1<sup>+</sup>::bqt4p-GFP-bqt4C<sup>414-432</sup>* | This study | 6A |
| TL250 | *h<sup>-</sup> bqt4Δ::hph lys1<sup>+</sup>::bqt4p-GFP-bqt4C<sup>369-425</sup>* | This study | 6A, C, S7 |
| TL251 | *h<sup>-</sup> bqt4Δ::hph bqt3Δ::NAT lys1<sup>+</sup>::bqt4p-GFP-bqt4C<sup>369-425</sup>* | This study | 6A |
| H1N546 | *h<sup>+</sup> bqt4Δ::hph lys1<sup>+</sup>::ish1-GFP* | This study | 7A |
| TL206 | *h<sup>-</sup> bqt4Δ::hph lys1<sup>+</sup>::nmt1p-GFP-bqt4* | This study | 7B-E |
| TL207 | *h<sup>-</sup> bqt4Δ::hph lys1<sup>+</sup>::nmt1p-GFP-bqt4 bqt3Δ::NAT* | This study | 7E |
| TL286 | *h<sup>-</sup> bqt4Δ::hph lys1<sup>+</sup>::bqt4p-GFP-bqt4 mts2-1* | This study | S1 |
| TL287 | *h<sup>-</sup> bqt4Δ::hph bqt3Δ::NAT lys1<sup>+</sup>::bqt4p-GFP-bqt4 mts2-1* | This study | S1 |
| TL222 | *h<sup>-</sup> bqt4Δ::hph bqt3Δ::NAT lys1<sup>+</sup>::bqt4p-GFP-bqt4 meu34Δ::kan<sup>r</sup>* | This study | S2 |
| TL224 | *h<sup>-</sup> bqt4Δ::hph bqt3Δ::NAT lys1<sup>+</sup>::bqt4p-GFP-bqt4 san1Δ::kan<sup>r</sup>* | This study | S2 |
| TL225 | *h<sup>-</sup> bqt4Δ::hph bqt3Δ::NAT lys1<sup>+</sup>::bqt4p-GFP-bqt4 ubr1Δ::kan<sup>r</sup>* | This study | S2 |
| TL226 | *h<sup>-</sup> bqt4Δ::hph bqt3Δ::NAT lys1<sup>+</sup>::bqt4p-GFP-bqt4 ubr11Δ::kan<sup>r</sup>* | This study | S2 |
| TL227 | *h<sup>-</sup> bqt4Δ::hph bqt3Δ::NAT lys1<sup>+</sup>::bqt4p-GFP-bqt4 hul5Δ::kan<sup>r</sup>* | This study | S2, S3A, B |
| TL267 | *h<sup>-</sup> bqt4Δ::hph bqt3Δ::NAT lys1<sup>+</sup>::bqt4p-GFP-bqt4 dsc1Δ::kan<sup>r</sup>* | This study | S2 |
| TL292 | *h<sup>-</sup> bqt4Δ::hph bqt3Δ::NAT lys1<sup>+</sup>::bqt4p-GFP-bqt4 cut9-665* | This study | S2 |
| TL294 | *h<sup>-</sup> bqt4Δ::hph bqt3Δ::NAT lys1<sup>+</sup>::bqt4p-GFP-bqt4 ptr1-1* | This study | S2 |
| TL235 | *h<sup>-</sup> bqt4Δ::hph bqt3Δ::NAT lys1<sup>+</sup>::bqt4p-GFP-bqt4 doa10Δ::bsd hul5Δ::kan<sup>r</sup>* | This study | S3A, B |
| TL262 | *h<sup>-</sup> lys1-131 bqt4Δ::hph lys1<sup>+</sup>::GFP-bqt4C<sup>369-425</sup> doa10Δ::bsd* | This study | S7 |

*S. cerevisiae* strains
| Strain | Genotype | Source | Figure |
| --- | --- | --- | --- |
| AH109 | *MATa, trp1-901, leu2-3, 112, ura3-52, his3-200, gal4Δ, gal80Δ, LYS2::GAL1<sub>UAS</sub>-GAL1<sub>TATA</sub>-HIS3, MEL1, GAL2<sub>UAS</sub>-GAL2<sub>TATA</sub>-ADE2, URA3::MEL1<sub>UAS</sub>-MEL1<sub>TATA</sub>-lacZ)* | Clontech, Takara Bio | 6B |
| H1N2190 | AH109 harboring pGADT7 and pGBKT7 | This study | 6B |
| H1N2193 | AH109 harboring pGADT7 and pGBKT7-bqt3 | This study | 6B |
| H1N2786 | AH109 harboring pGADT7-GFP-bqt4(1-432) and pGBKT7 | This study | 6B |
| H1N2787 | AH109 harboring pGADT7-GFP-bqt4(1-432) and pGBKT7-bqt3 | This study | 6B |
| H1N2788 | AH109 harboring pGADT7-GFP-bqt4(369-432) and pGBKT7-bqt3 | This study | 6B |
| H1N2789 | AH109 harboring pGADT7-GFP-bqt4(394-432) and pGBKT7-bqt3 | This study | 6B |
| H1N2790 | AH109 harboring pGADT7-GFP-bqt4(414-432) and pGBKT7-bqt3 | This study | 6B |
| H1N2791 | AH109 harboring pGADT7-GFP-bqt4(1-413) and pGBKT7-bqt3 | This study | 6B |
| H1N2792 | AH109 harboring pGADT7-GFP-bqt4(1-425) and pGBKT7-bqt3 | This study | 6B |
| H1N2793 | AH109 harboring pGADT7-GFP-bqt4(369-425) and pGBKT7-bqt3 | This study | 6B |
| H1N2794 | AH109 harboring pGADT7-GFP-bqt4(414-425) and pGBKT7-bqt3 | This study | 6B |
| H1N2795 | AH109 harboring pGADT7-GFP-bqt4(426-432) and pGBKT7-bqt3 | This study | 6B |

**Supplementary Fig. 1.**
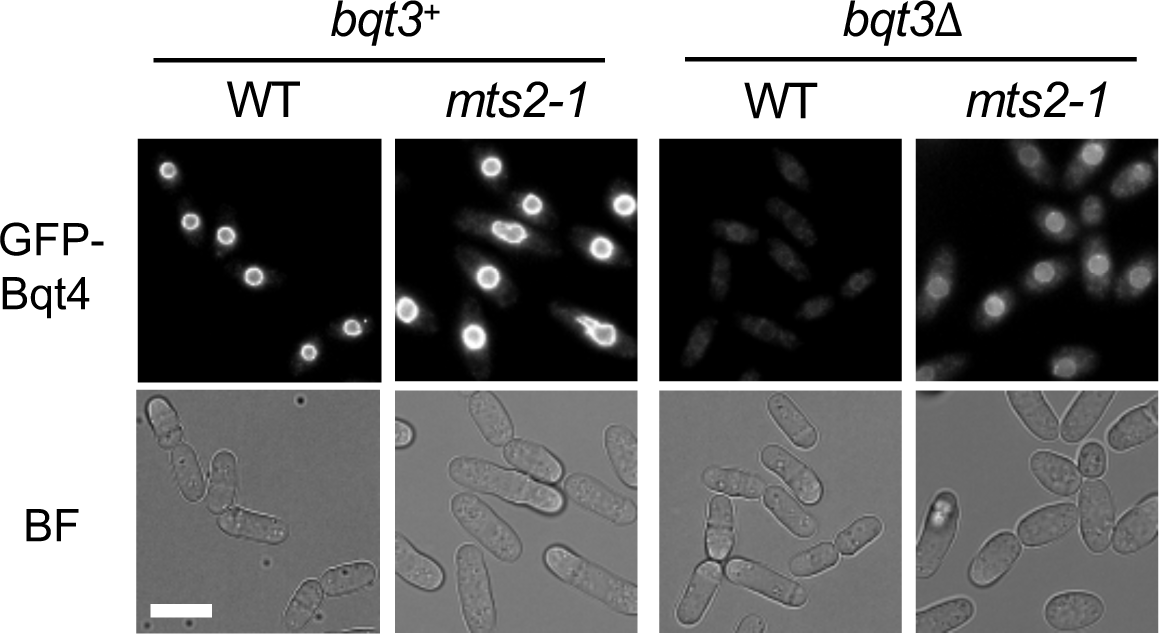
The *mts2-1* mutation prevents Bqt4 degradation. Cells harboring *mts2-1* in the *bqt3*^+^ or *bqt3*Δ background were cultured at a nonpermissive temperature (36°C) for 4 h and observed by fluorescence microscopy. (Upper panels) Fluorescence images of GFP-Bqt4 (lower panels) and bright-field images (BF). Bar: 10 μm.

**Supplementary Fig. 2.**
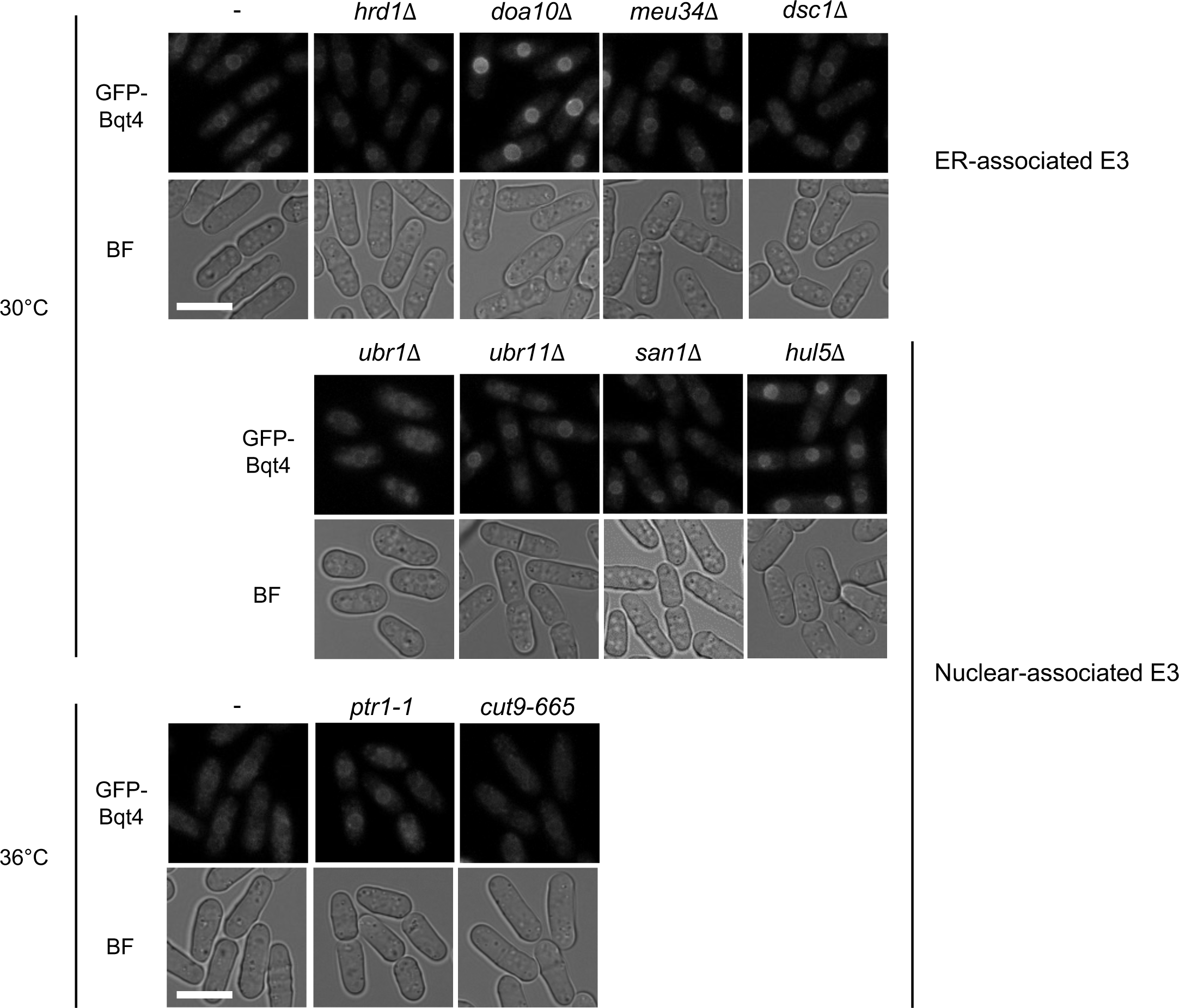
Most nuclear- or ER-associated E3 ligases have a minimal role in Bqt4 degradation. Fluorescence images of GFP-Bqt4 in the mutants are shown. Deletion or mutation of E3 ligases localized in the ER (*meu34*τι, *dsc1*τι) or functioning in the nucleus (*san1*τι, *ubr1*τι, *ubr11*τι, *hul5*τι, *ptr1-1*, *cut9-665*) was introduced in *bqt3*Δ cells expressing GFP-Bqt4. The cells were cultured at 30°C for 16 h or shifted to 36°C for 4 h for *ts* mutants and then observed using fluorescence microscopy. Bars: 10 μm.

**Supplementary Fig. 3.**
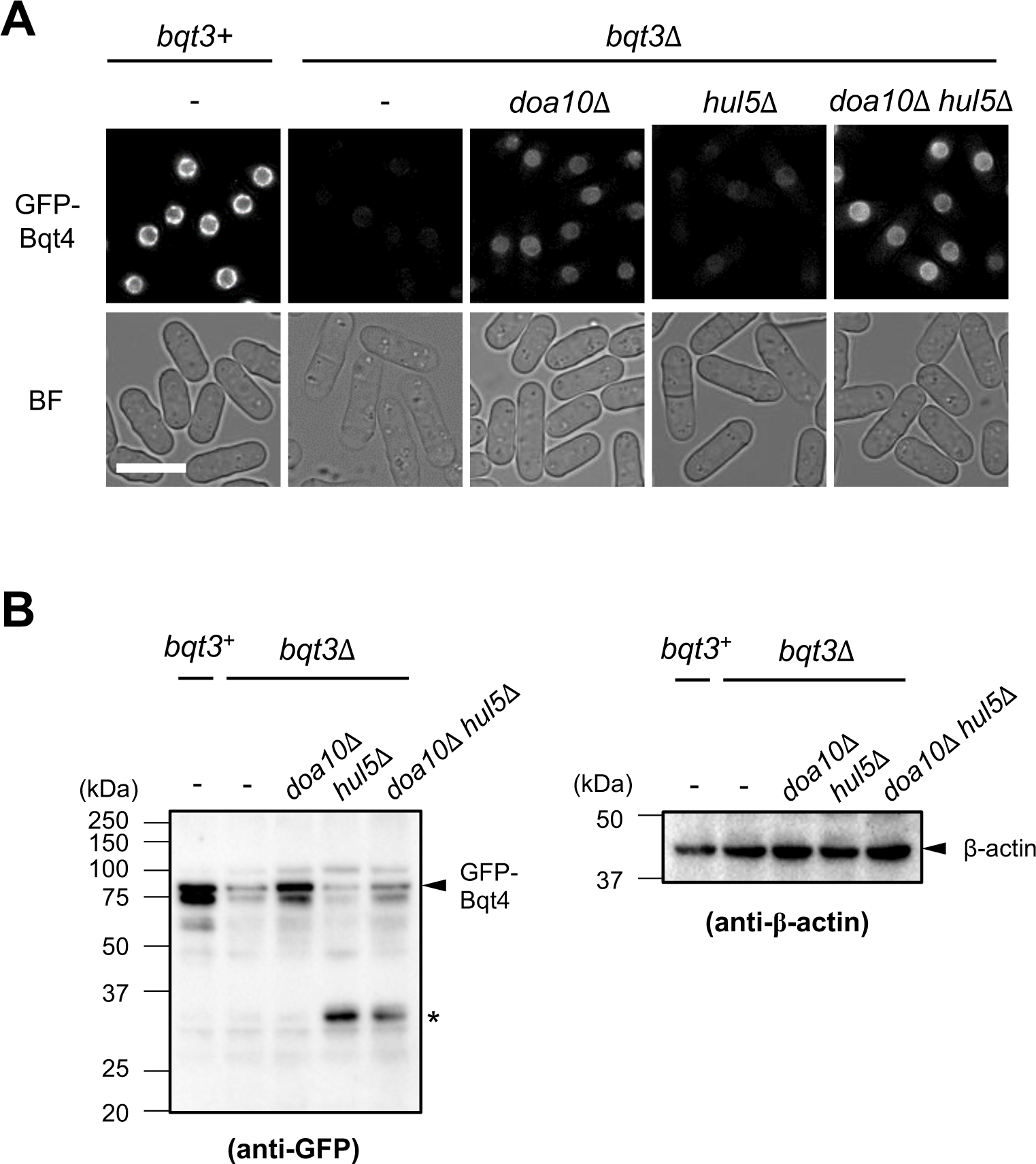
Synthetic effect of double deletion of *doa10* and *hul5* for GFP-Bqt4 level. Cells harboring the single (*doa10*11, *hul5*11) or double (*doa10*11*hul5*11) deletion of ubiquitin E3 ligase in *bqt3*Δ background were cultured at 30°C and subjected to microscopic observation (A) or western blotting (B). (A) Fluorescent images of GFP-Bqt4 mutants are shown. “*bqt3*^+^” shows GFP-Bqt4 level in *bqt3*^+^ cells. BF represents the bright-field image. Bar: 10 μm. (B) Protein levels of GFP-Bqt4 and β-actin were detected by anti-GFP and anti-β-actin antibodies, respectively. The molecular weight markers are shown on the left. The asterisk indicates the degraded product.

**Supplementary Fig. 4.**
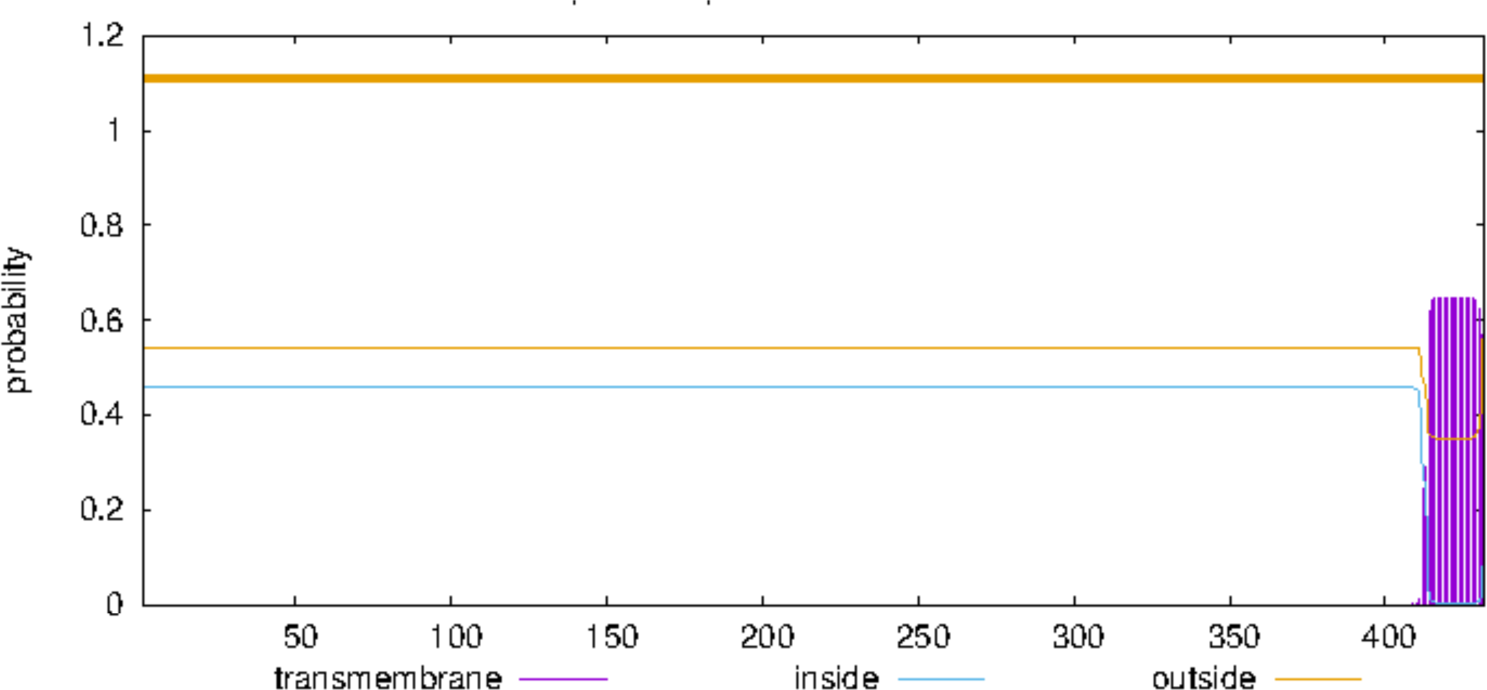
Prediction of Bqt4 TMD properties. The transmembrane helices of Bqt4 were predicted using the TMHMM-2.0 prediction server (https://services.healthtech.dtu.dk/service.php?TMHMM-2.0). The horizontal and vertical axes indicate the prediction probability and amino acid residues of Bqt4, respectively.

**Supplementary Fig. 5.**
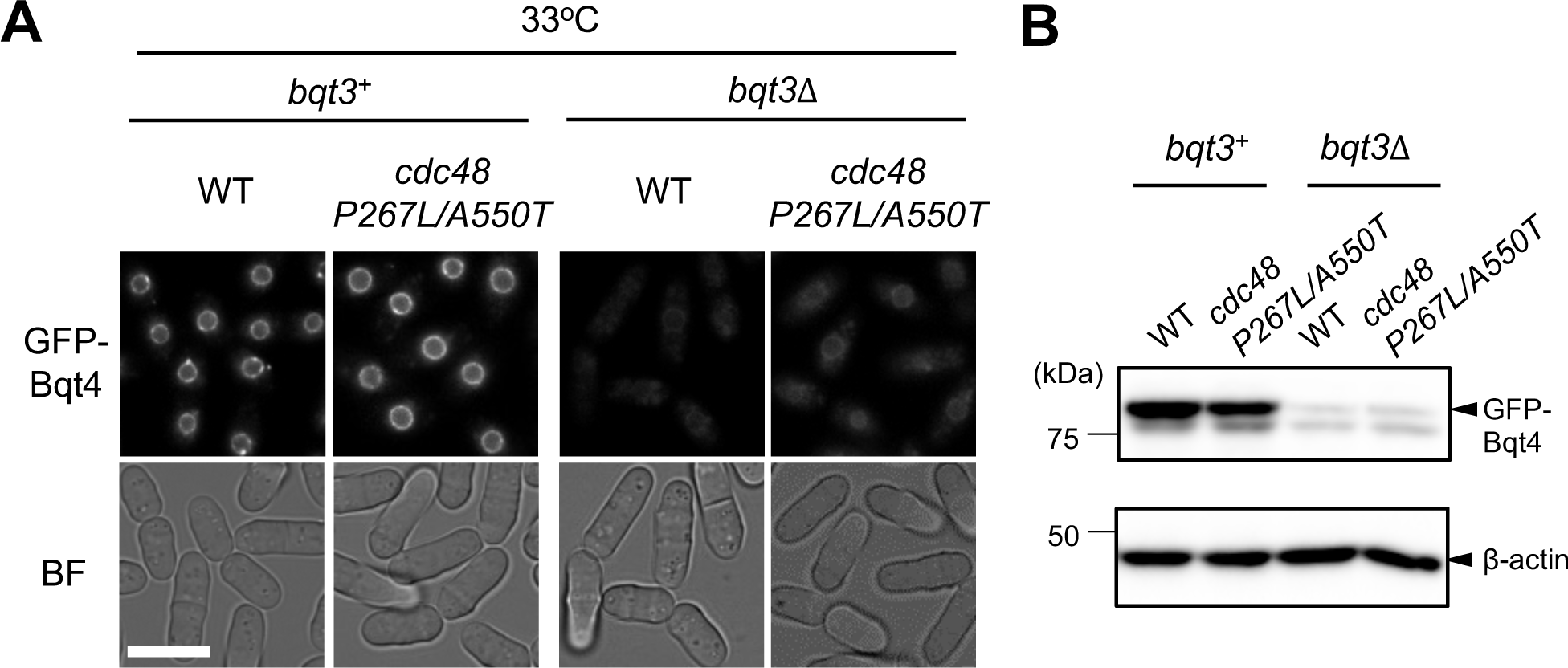
Degradation of Bqt4 is not significantly affected in Cdc48 mutant at the permissive temperature. Cells harboring the cdc48-P267L/A550T mutant in the *bqt3+* or *bqt3Δ* background were cultured at 33°C for 15 h and subjected to microscopic observation (A) or western blotting (B). (A) Fluorescence images of GFP-Bqt4 (upper panels) and bright-field images (lower panels). Bar: 10 μm. (B) The protein levels of GFP-Bqt4 and β-actin were detected by anti-GFP and anti-β-actin antibodies, respectively. The molecular weight markers are shown on the left.

**Supplementary Fig. 6.**
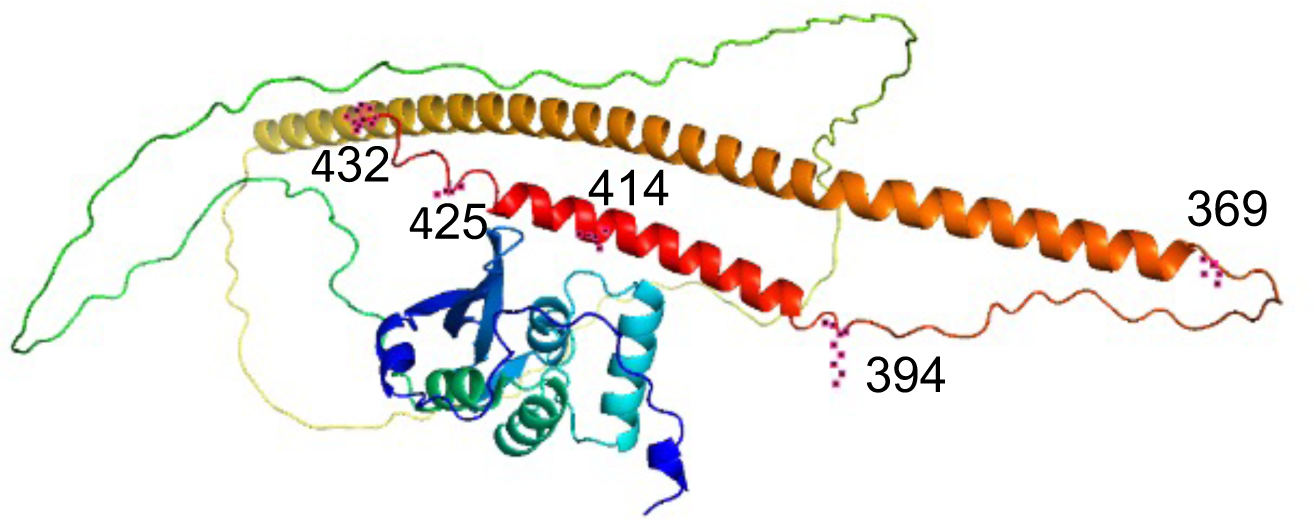
Structure of Bqt4 as predicted by AlphaFold2. The three-dimensional structure of Bqt4 predicted using AlphaFold2 was obtained from the website (https://alphafold.ebi.ac.uk/entry/O60158).

**Supplementary Fig. 7.**
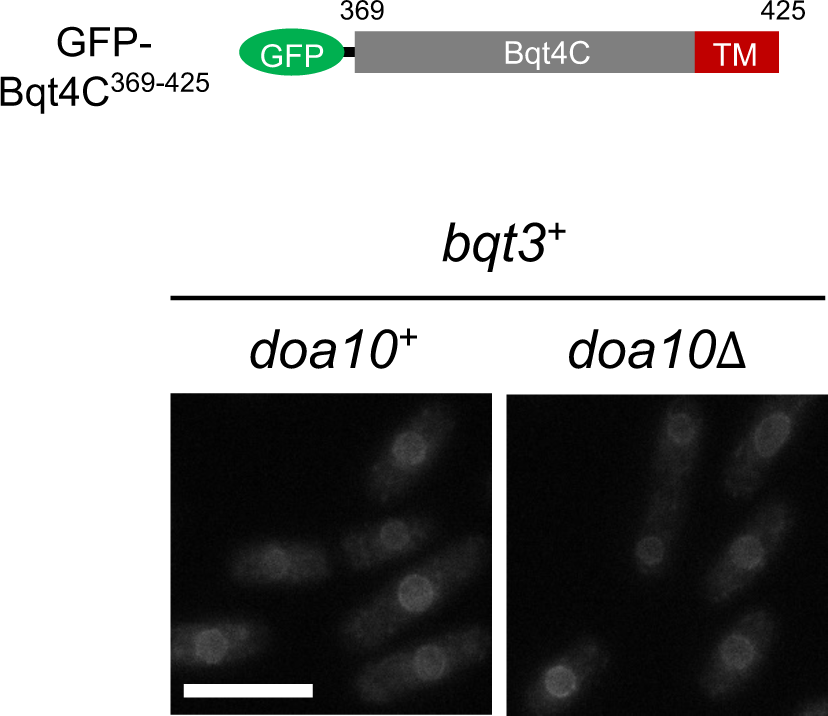
Degradation of Bqt4C^369-425^ occurred independently of Doa10 pathway. The GFP-Bqt4C^369-425^ fragment was expressed in *doa10*^+^ and *doa10*Δ cells and observed by fluorescence microscopy. Scale bar: 10 μm.

